## Supplementary material for "Large-Scale Statistical Dissection of Sequence-Derived Biochemical Features Distinguishing Soluble and Insoluble Proteins": https://github.com/huyhoang6723/protein-solubility-effectsize

### Supplementary Tables

Table 1: **Table S1. Comprehensive statistical results for 16 global and grouped-composition sequence-derived descriptors.**

| **Feature** | **Soluble median [Q1, Q3]** | **Insoluble median [Q1, Q3]** | **HL [95% CI]** | **$\delta$ [95% CI]** | **FDR $q$** | **AUC** | $J$ | $T^*$ |
|---|---|---|---|---|---|---|---|---|
| length | 209 [129, 338] | 277 [172, 426] | -70 [-77, -62] | -0.215 [-0.237, -0.196] | 0.000 | 0.392 | 0.158 | 227 |
| molecular_weight | 19 743 [12 201, 31 715] | 25 966 [16 148, 39 869] | -6322.485 [-7147.547, -5784.646] | -0.214 [-0.244, -0.204] | 0.000 | 0.393 | 0.157 | 21520 |
| aggregation_ratio | 0.02262 [0.01502, 0.03371] | 0.01869 [0.01252, 0.02857] | 0.004 [0.003, 0.004] | 0.166 [0.138, 0.178] | 0.000 | 0.583 | 0.125 | 0.020 |
| neg_ratio | 0.1298 [0.1104, 0.1510] | 0.1219 [0.1034, 0.1414] | 0.008 [0.007, 0.009] | 0.150 [0.134, 0.173] | 0.000 | 0.575 | 0.114 | 0.126 |
| isoelectric_point | 6.401 [4.971, 8.357] | 6.931 [5.384, 8.941] | -0.588 [-0.702, -0.468] | -0.132 [-0.167, -0.125] | 0.000 | 0.434 | 0.102 | 6.523 |
| net_charge_pH7 | -1.882 [-7.066, 3.770] | -0.207 [-5.000, 6.476] | -1.618 [-1.931, -1.217] | -0.120 [-0.143, -0.099] | 0.000 | 0.440 | 0.107 | 2.413 |
| mean_hydropathy | -0.3511 [-0.4695, -0.2277] | -0.3085 [-0.4388, -0.1834] | -0.046 [-0.060, -0.035] | -0.080 [-0.106, -0.064] | 0.000 | 0.460 | 0.067 | -0.305 |
| sulfur_ratio | 0.03448 [0.01961, 0.04712] | 0.03681 [0.02193, 0.05013] | -0.002 [-0.003, -0.002] | -0.076 [-0.096, -0.056] | 0.000 | 0.462 | 0.066 | 0.029 |
| helix_prop_mean | 1.029 [1.019, 1.038] | 1.025 [1.015, 1.034] | 0.004 [0.003, 0.005] | 0.060 [0.049, 0.089] | 0.000 | 0.530 | 0.054 | 1.041 |
| hydrophobic_ratio | 0.4019 [0.3786, 0.4253] | 0.4063 [0.3829, 0.4286] | -0.003 [-0.005, -0.001] | -0.051 [-0.053, -0.012] | 0.000 | 0.475 | 0.046 | 0.419 |
| sheet_prop_mean | 0.9937 [0.9857, 1.0024] | 0.9963 [0.9883, 1.0048] | -0.002 [-0.003, -0.000] | -0.050 [-0.056, -0.015] | 0.000 | 0.475 | 0.046 | 1.008 |
| disorder_ratio | 0.3976 [0.3690, 0.4310] | 0.3937 [0.3662, 0.4274] | 0.003 [0.001, 0.005] | 0.048 [0.021, 0.063] | 0.000 | 0.524 | 0.039 | 0.412 |
| polar_ratio | 0.2006 [0.1765, 0.2247] | 0.2031 [0.1798, 0.2278] | -0.002 [-0.005, -0.001] | -0.024 [-0.055, -0.013] | 0.000 | 0.488 | 0.023 | 0.195 |
| tiny_ratio | 0.1389 [0.1111, 0.1667] | 0.1411 [0.1127, 0.1695] | -0.000 [-0.003, 0.002] | -0.021 [-0.035, 0.009] | 0.000 | 0.489 | 0.037 | 0.171 |
| pos_ratio | 0.1412 [0.1212, 0.1636] | 0.1419 [0.1230, 0.1636] | -0.000 [-0.002, 0.001] | -0.015 [-0.026, 0.015] | 0.000 | 0.492 | 0.022 | 0.116 |
| turn_prop_mean | 0.9667 [0.9584, 0.9759] | 0.9672 [0.9589, 0.9763] | -0.001 [-0.003, 0.000] | -0.011 [-0.036, 0.004] | 0.011 | 0.495 | 0.021 | 0.928 |

**Interpretive note.** This table reports the full statistical summary for 16 global and grouped-composition descriptors used to compare soluble and insoluble proteins. The largest absolute effects are observed for `length` and `molecular_weight`, both with negative Cliff's $\delta$ values ($-0.215$ and $-0.214$), indicating that insoluble proteins tend to be longer and heavier. The corresponding Hodges–Lehmann shifts indicate a median displacement of 70 residues and 6322.485 Da toward the insoluble class.

Among positively oriented descriptors, `aggregation_ratio` ($\delta = 0.166$, AUC $= 0.583$) and `neg_ratio` ($\delta = 0.150$, AUC $= 0.575$) show the strongest enrichment in soluble proteins. Their effect sizes are statistically robust but remain modest in magnitude, consistent with weak but coherent discriminatory structure rather than strong class separation.

Charge-related descriptors such as `isoelectric_point` and `net_charge_pH7` exhibit intermediate negative effects, suggesting that insoluble proteins tend to have higher pI values and reduced net negativity. Hydropathy- and sulfur-related descriptors also show systematic but small shifts, whereas disorder and secondary-structure propensity features remain close to the null regime.

Overall, the combination of very small FDR-adjusted $q$-values with mostly modest $|\delta|$, low Youden's $J$, and AUC values near 0.5 indicates that statistical significance is widespread, but practical univariate discrimination is limited. The table therefore supports a weak-signal interpretation in which multiple correlated and partially overlapping physicochemical features contribute jointly to solubility differences.

Table 2: **Table S2. Comprehensive statistical results for amino acid frequency descriptors.**

| **Feature** | **Soluble median [Q1, Q3]** | **Insoluble median [Q1, Q3]** | **HL [95% CI]** | $\boldsymbol{\delta}$ **[95% CI]** | **FDR** $q$ | **AUC** | $J$ | $T^*$ |
|---|---|---|---|---|---|---|---|---|
| freq_C | 0.01017 [0.00312, 0.01882] | 0.01282 [0.00615, 0.02143] | -0.003 [-0.003, -0.002] | -0.131 [-0.151, -0.111] | 0.000 | 0.434 | 0.100 | 0.000 |
| freq_E | 0.07064 [0.05488, 0.08889] | 0.06529 [0.05072, 0.08187] | 0.006 [0.005, 0.007] | 0.121 [0.119, 0.160] | 0.000 | 0.560 | 0.092 | 0.071 |
| freq_R | 0.05350 [0.03663, 0.07273] | 0.05894 [0.04061, 0.07944] | -0.005 [-0.006, -0.005] | -0.105 [-0.101, -0.061] | 0.000 | 0.447 | 0.083 | 0.060 |
| freq_K | 0.06019 [0.04255, 0.07843] | 0.05350 [0.03636, 0.07339] | 0.007 [0.006, 0.008] | 0.091 [0.067, 0.109] | 0.000 | 0.546 | 0.080 | 0.049 |
| freq_D | 0.05721 [0.04098, 0.07391] | 0.05473 [0.04000, 0.07074] | 0.003 [0.002, 0.003] | 0.080 [0.055, 0.097] | 0.000 | 0.540 | 0.067 | 0.061 |
| freq_S | 0.06349 [0.04724, 0.08065] | 0.06630 [0.05000, 0.08333] | -0.003 [-0.003, -0.002] | -0.068 [-0.092, -0.044] | 0.000 | 0.466 | 0.043 | 0.067 |
| freq_P | 0.04244 [0.02899, 0.05755] | 0.04494 [0.03125, 0.06061] | -0.003 [-0.003, -0.002] | -0.064 [-0.088, -0.040] | 0.000 | 0.468 | 0.050 | 0.048 |
| freq_A | 0.07273 [0.05556, 0.09091] | 0.07650 [0.05825, 0.09524] | -0.004 [-0.005, -0.003] | -0.051 [-0.075, -0.026] | 0.000 | 0.474 | 0.052 | 0.092 |
| freq_Q | 0.03768 [0.02500, 0.05172] | 0.03643 [0.02424, 0.05000] | 0.001 [0.001, 0.002] | 0.042 [0.017, 0.066] | 0.000 | 0.521 | 0.042 | 0.030 |
| freq_Y | 0.03061 [0.01923, 0.04211] | 0.02913 [0.01818, 0.04110] | 0.001 [0.001, 0.002] | 0.041 [0.016, 0.066] | 0.000 | 0.521 | 0.041 | 0.009 |
| freq_W | 0.01000 [0.00000, 0.01754] | 0.01031 [0.00000, 0.01852] | -0.000 [-0.001, 0.000] | -0.039 [-0.063, -0.014] | 0.000 | 0.481 | 0.047 | 0.000 |
| freq_I | 0.05128 [0.03604, 0.06818] | 0.05333 [0.03788, 0.07059] | -0.002 [-0.003, -0.001] | -0.039 [-0.063, -0.014] | 0.000 | 0.481 | 0.047 | 0.045 |
| freq_V | 0.06140 [0.04545, 0.07813] | 0.06349 [0.04688, 0.08000] | -0.002 [-0.003, -0.001] | -0.035 [-0.060, -0.010] | 0.000 | 0.482 | 0.043 | 0.077 |
| freq_N | 0.03922 [0.02632, 0.05263] | 0.04000 [0.02740, 0.05405] | -0.001 [-0.002, -0.000] | -0.033 [-0.058, -0.008] | 0.000 | 0.484 | 0.040 | 0.056 |
| freq_G | 0.06195 [0.04598, 0.07937] | 0.06383 [0.04688, 0.08108] | -0.002 [-0.003, -0.001] | -0.028 [-0.053, -0.003] | 0.000 | 0.487 | 0.039 | 0.031 |
| freq_L | 0.09091 [0.07317, 0.10976] | 0.09195 [0.07407, 0.11111] | -0.001 [-0.002, -0.000] | -0.026 [-0.051, -0.001] | 0.000 | 0.487 | 0.038 | 0.079 |
| freq_F | 0.03636 [0.02532, 0.04878] | 0.03704 [0.02632, 0.05000] | -0.001 [-0.001, -0.000] | -0.018 [-0.043, 0.007] | 0.000 | 0.491 | 0.029 | 0.018 |
| freq_H | 0.02381 [0.01389, 0.03448] | 0.02439 [0.01471, 0.03571] | -0.001 [-0.001, 0.000] | -0.016 [-0.041, 0.009] | 0.000 | 0.492 | 0.029 | 0.012 |
| freq_M | 0.02174 [0.01220, 0.03125] | 0.02174 [0.01282, 0.03125] | 0 [-0.001, 0.001] | -0.003 [-0.028, 0.023] | 0.514 | 0.499 | 0.023 | 0.011 |
| freq_T | 0.05263 [0.03704, 0.06897] | 0.05263 [0.03788, 0.06897] | 0 [-0.001, 0.001] | 0.000 [-0.025, 0.025] | 0.989 | 0.500 | 0.022 | 0.072 |

**Interpretive note.** This table summarizes the statistical comparison of individual amino acid frequencies between soluble and insoluble proteins. Compared to global descriptors (Table S1), all amino acid-level effects are substantially smaller in magnitude, indicating that no single residue frequency provides strong standalone discrimination.

The largest effects are observed for charged residues. Glutamate (`freq_E`) shows a positive effect ($\delta = 0.121$, AUC $= 0.560$), indicating enrichment in soluble proteins, whereas arginine (`freq_R`) and cysteine (`freq_C`) exhibit negative effects ($\delta = -0.105$ and $-0.131$), suggesting higher prevalence in insoluble proteins. Lysine (`freq_K`) and aspartate (`freq_D`) show moderate positive effects, reinforcing the role of electrostatic composition in solubility modulation.

Hydrophobic and aliphatic residues (e.g., `freq_I`, `freq_V`, `freq_L`) exhibit weak negative effects, consistent with slightly higher abundance in insoluble proteins, but with small effect sizes ($|\delta| < 0.05$) and AUC values close to 0.5. Similarly, most polar and small residues show negligible differences.

Notably, methionine (`freq_M`) and threonine (`freq_T`) are not statistically significant after FDR correction ($q = 0.514$ and $0.989$), indicating no detectable distributional difference.

Overall, although many amino acid frequencies are statistically significant due to large sample size, their small effect sizes, low Youden's $J$, and AUC values near 0.5 demonstrate that individual residue composition alone provides limited discriminative power. These results support a weak-signal interpretation in which solubility differences arise from the collective contribution of multiple features rather than dominant effects from single amino acids.

Table 3: **Table S3. Robust summary statistics defining the two orthogonal physicochemical axes of the composite-$\delta$ index.**

| **Feature** | **Median** | **IQR** |
|---|---|---|
| length | 236 | 231 |
| neg_ratio | 0.126531 | 0.039934 |

**Interpretive note.** This table reports the robust summary statistics used for centering and scaling in the redundancy-aware composite-$\delta$ formulation. The selected features correspond to two approximately orthogonal physicochemical dimensions identified after redundancy filtering: a structural burden axis represented by sequence length, and an electrostatic stabilization axis represented by the proportion of negatively charged residues.

The median values define the central tendency of each feature across the combined dataset, while the interquartile range (IQR) provides a robust measure of dispersion that is insensitive to heavy-tailed distributions and outliers. Notably, sequence length exhibits a very large IQR (231 residues), reflecting substantial variability across proteins, whereas the negative charge proportion shows a much narrower spread (IQR $\approx 0.04$), indicating tighter distributional concentration.

These contrasting scales highlight the need for robust normalization when combining features of different magnitudes. By centering at the median and scaling by IQR, the composite-$\delta$ index ensures that both structural and electrostatic contributions are comparably weighted without being dominated by scale differences.

Table 4: **Table S4. Comparative performance of reference solubility predictors and the composite-$\delta$ baseline.**

| Model | $T$ | AUC | Acc. | F1 | MCC | Prec. | Sens. | Spec. |
|---|---|---|---|---|---|---|---|---|
| Protein_sol | 0.5 | 0.5985 | 0.5649 | 0.6350 | 0.1405 | 0.5470 | 0.7569 | 0.3729 |
| SKADE | 0.5 | 0.6882 | 0.6443 | 0.5304 | 0.3305 | 0.7811 | 0.4015 | 0.8874 |
| SWI | 0.5 | 0.5597 | 0.5423 | 0.6334 | 0.0971 | 0.5284 | 0.7905 | 0.2938 |
| Solupro | 0.5 | 0.7126 | 0.6308 | 0.6639 | 0.2667 | 0.6095 | 0.7290 | 0.5325 |
| EPSOL | 0.5 | 0.6664 | 0.6663 | 0.5915 | 0.3578 | 0.7630 | 0.4830 | 0.8499 |
| NetSolP | 0.5 | 0.6183 | 0.5502 | 0.6557 | 0.1268 | 0.5312 | 0.8564 | 0.2437 |
| DeepSol-E | 0.4 | 0.6660 | 0.5990 | 0.6407 | 0.2036 | 0.5804 | 0.7150 | 0.4830 |
| PLM_Sol | 0.5 | 0.8342 | 0.7299 | 0.7542 | 0.4690 | 0.6919 | 0.8289 | 0.6308 |
| **Composite-$\delta$** | 0 | **0.6240** | **0.5734** | **0.5875** | **0.1746** | **0.6922** | **0.5102** | **0.6663** |

**Interpretive note.** This table provides a contextual performance comparison between the proposed composite-$\delta$ baseline and a range of established solubility prediction models. High-capacity approaches, particularly PLM-based models, achieve substantially higher predictive performance (AUC = 0.8342, MCC = 0.4690), reflecting their ability to capture higher-order sequence dependencies and contextual information.

In contrast, the composite-$\delta$ index achieves moderate performance (AUC = 0.6240, MCC = 0.1746), comparable to or exceeding several classical feature-based models (e.g., Protein_sol, SWI). Importantly, this level of performance is achieved without any model training, parameter optimization, or sequence embedding, relying solely on two global sequence-derived descriptors.

A notable characteristic of the composite-$\delta$ model is its balanced behavior, with relatively high precision (0.6922) and specificity (0.6663), but lower sensitivity (0.5102), indicating a tendency toward conservative predictions of solubility. This contrasts with several existing models that favor high sensitivity at the expense of specificity.

Overall, the comparison highlights a fundamental trade-off: while deep learning models provide superior predictive accuracy, the composite-$\delta$ baseline offers a transparent, computationally efficient, and interpretable alternative. It serves as a lower-bound reference for the discriminative information contained in global physicochemical features and provides a practical tool for rapid preliminary screening in resource-constrained settings.

Table 5: **Table S5. Computational complexity comparison across solubility prediction models.**

| **Model** | **Architecture Type** | **Inference Complexity** | **Training Requirement** |
|---|---|---|---|
| Protein_sol | Feature-based ML | $O(d)$–$O(t)$ | Supervised training |
| SKADE | Ensemble model | $O(t)$ | Supervised training |
| SWI | Linear sequence index | $O(L)$ | None |
| Solupro | Feature-based ML | $O(d)$–$O(t)$ | Supervised training |
| EPSOL | Ensemble / ML | $O(t)$ | Supervised training |
| NetSolP | Deep neural network | $O(L \cdot k)$ | Supervised training |
| DeepSol-E | CNN-based model | $O(L \cdot k)$ | Supervised training |
| PLM_Sol | Transformer (PLM-based) | $O(H \cdot L^2 \cdot h)$ | Pretrained + fine-tuned |
| **Composite-$\delta$** | Interpretable linear score | $\mathbf{O(L)}$ (feature extraction) + $\mathbf{O(1)}$ (scoring) | None |

**Interpretive note.** This table provides a qualitative comparison of computational complexity across representative solubility prediction models. The methods span a wide range of architectural paradigms, from simple interpretable scoring schemes to deep neural networks and transformer-based protein language models.

Feature-based and ensemble methods typically operate with complexity dependent on the number of extracted descriptors ($O(d)$) or ensemble size ($O(t)$), whereas sequence-based neural architectures scale linearly with sequence length ($O(L \cdot k)$). Transformer-based protein language models exhibit substantially higher computational cost, scaling quadratically with sequence length ($O(H \cdot L^2 \cdot h)$), reflecting attention mechanisms over all residue pairs.

The composite-$\delta$ baseline differs fundamentally in that it does not require model training and involves only two global sequence-derived features. Its computational cost is dominated by feature extraction, which scales linearly with sequence length ($O(L)$), followed by constant-time scoring ($O(1)$). This results in negligible computational overhead compared to deep learning models.

Overall, the comparison highlights a clear trade-off between predictive performance and computational cost. While high-capacity models achieve superior accuracy, the composite-$\delta$ approach offers a lightweight, interpretable, and training-free alternative suitable for rapid large-scale screening and resource-constrained environments.

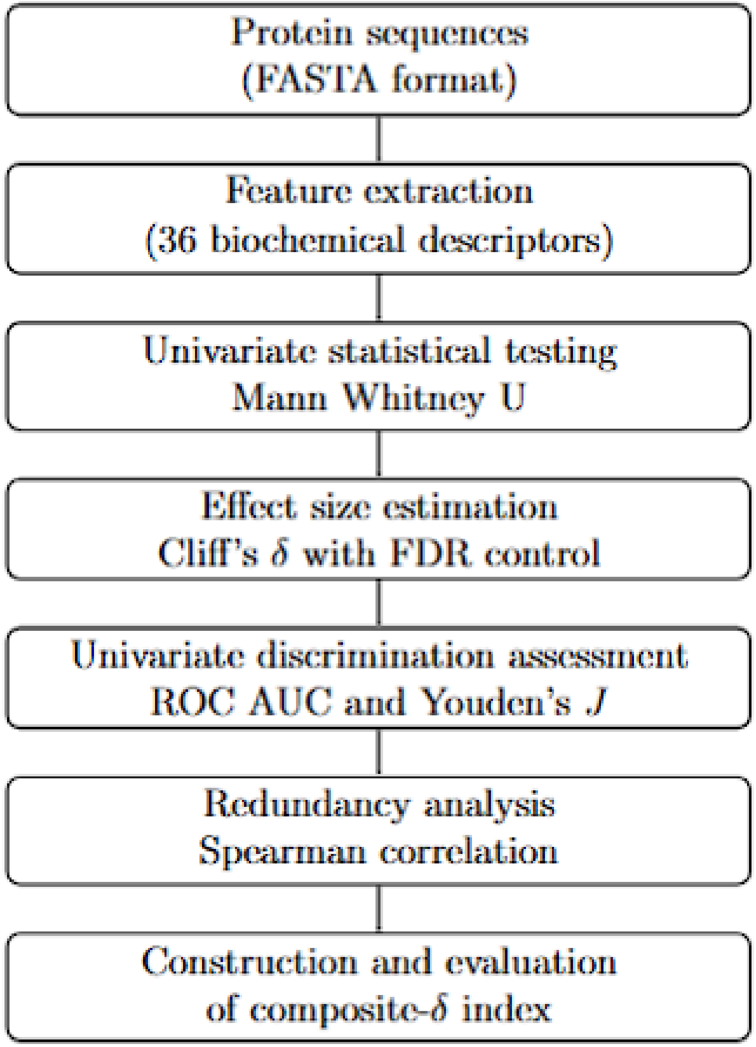


Table 6: **Figure S1. Analytical workflow of the proposed framework.** The diagram illustrates the complete analysis pipeline, including dataset acquisition, preprocessing of protein sequences, extraction of sequence-derived biochemical descriptors, and subsequent statistical analysis. The workflow comprises non-parametric hypothesis testing (Mann–Whitney U test), effect size estimation (Cliff's $\delta$), confidence interval estimation (Hodges–Lehmann), redundancy assessment via Spearman correlation, and construction of the composite-$\delta$ index. This pipeline emphasizes interpretability, statistical rigor, and reproducibility without reliance on model training.

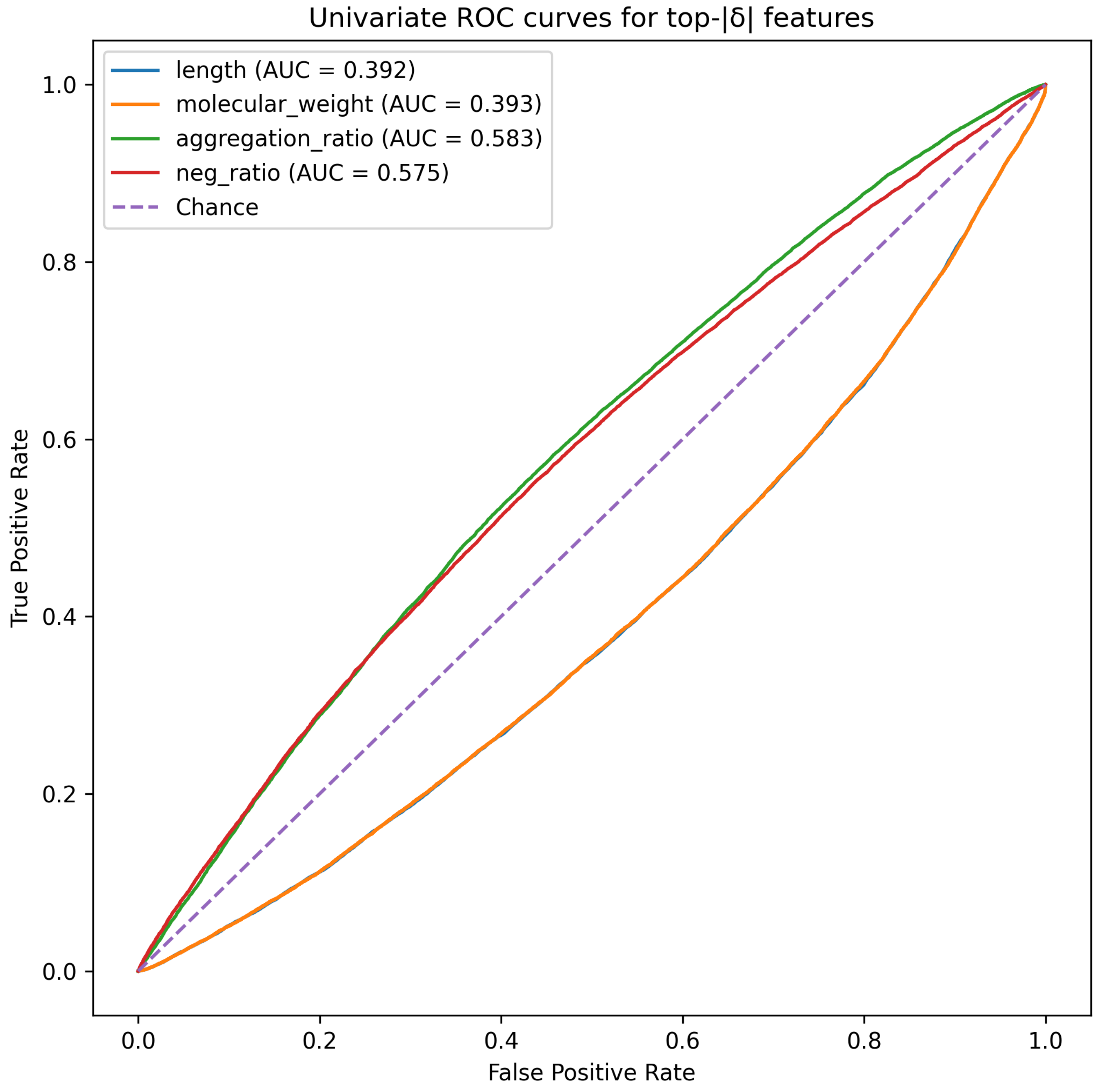


Table 7: **Figure S2. Univariate ROC curves for the four features with the largest absolute effect sizes.** Receiver operating characteristic (ROC) curves are shown for the top-ranked features based on absolute Cliff's $\delta$. Despite statistical significance, all curves remain close to the diagonal, indicating limited discriminative power at the individual feature level. This observation reinforces the weak-signal regime, where no single descriptor provides strong standalone classification performance.

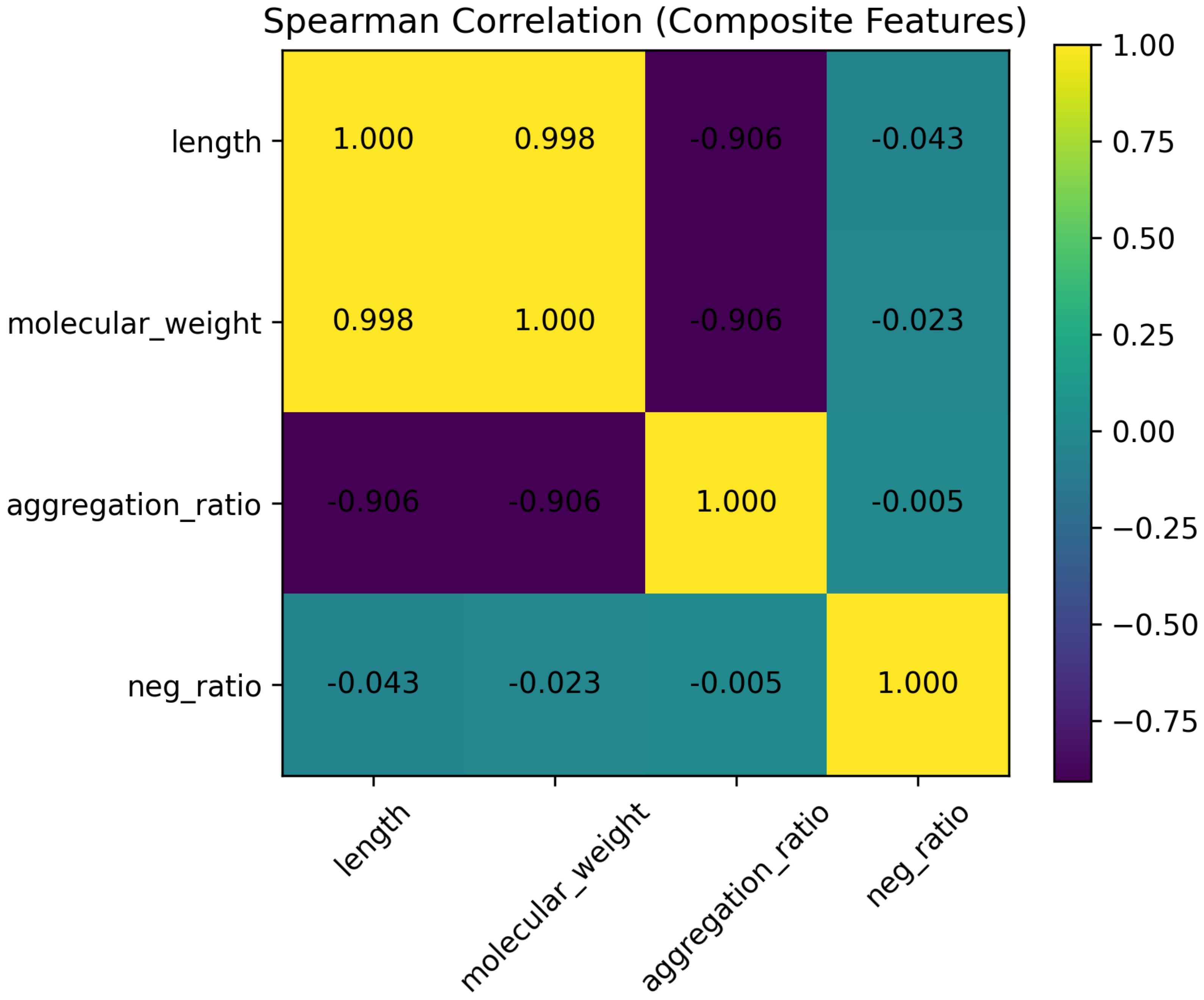


Table 8: **Figure S3. Spearman correlation matrix among top-ranked global descriptors.** The matrix illustrates pairwise monotonic relationships between selected physicochemical features. Strong correlations are observed among size-related descriptors (e.g., sequence length and molecular weight), indicating redundancy and the presence of a shared latent structural dimension. In contrast, electrostatic features such as negative charge proportion remain largely independent. Feature pairs with $|\rho| \geq 0.85$ were considered redundant and excluded in the construction of the composite-$\delta$ index.

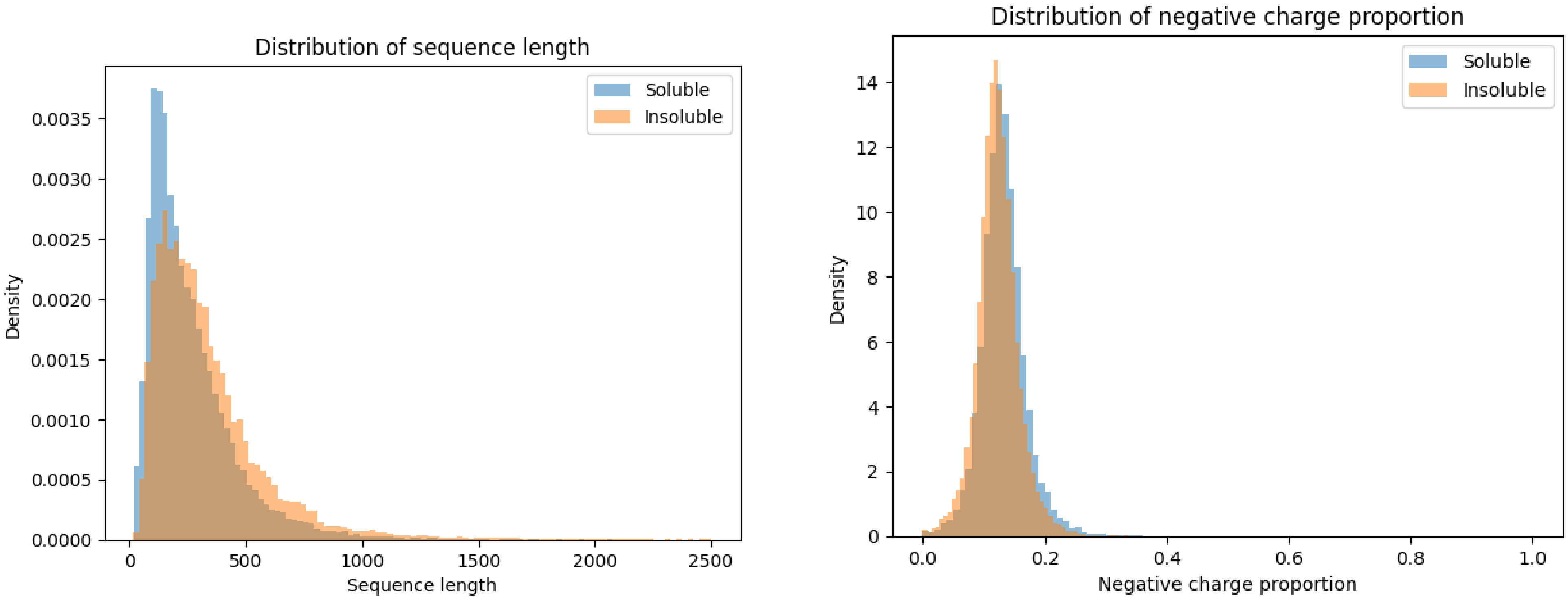


Table 9: **Figure S4. Distributional overlap of the two principal orthogonal features.** Kernel density estimates are shown for sequence length (left) and negative charge proportion (right) across soluble and insoluble proteins. Although statistically significant differences exist, both features exhibit substantial overlap between classes. Insoluble proteins tend to be longer, while soluble proteins show modest enrichment in negatively charged residues; however, these shifts are small relative to within-class variability. The extensive overlap highlights the intrinsic difficulty of classification using individual descriptors and supports the interpretation of protein solubility as a weak-signal phenomenon at the sequence level.
