## Supplementary material for "Large-Scale Statistical Dissection of Sequence-Derived Biochemical Features Distinguishing Soluble and Insoluble Proteins": https://github.com/huyhoang6723/protein-solubility-effectsize: main.tex

% Template for PLoS
% Version 3.7 Aug 2025
%
% % % % % % % % % % % % % % % % % % % % % %
%
% -- IMPORTANT NOTE
%
% This template contains comments intended 
% to minimize problems and delays during our production 
% process. Please follow the template instructions
% whenever possible.
%
% % % % % % % % % % % % % % % % % % % % % % % 
%
% Once your paper is accepted for publication, 
% PLEASE REMOVE ALL TRACKED CHANGES in this file 
% and leave only the final text of your manuscript. 
% PLOS recommends the use of latexdiff to track changes during review, as this will help to maintain a clean tex file.
% Visit https://www.ctan.org/pkg/latexdiff?lang=en for info or contact us at.
%
%
% There are no restrictions on package use within the LaTeX files except that no packages listed in the template may be deleted.
%
% Please do not include colors or graphics in the text.
%
% The manuscript LaTeX source should be contained within a single file (do not use \input, \externaldocument, or similar commands).
%
% % % % % % % % % % % % % % % % % % % % % % %
%
% -- FIGURES AND TABLES
%
% Please include tables/figure captions directly after the paragraph where they are first cited in the text.
%
% DO NOT INCLUDE GRAPHICS IN YOUR MANUSCRIPT
% - Figures should be uploaded separately from your manuscript file. 
% - Figures generated using LaTeX should be extracted and removed from the PDF before submission. 
% - Figures containing multiple panels/subfigures must be combined into one image file before submission.
% For figure citations, please use "Fig" instead of "Figure".
% See http://journals.plos.org/plosone/s/figures for PLOS figure guidelines.
%
% Tables should be cell-based and may not contain:
% - spacing/line breaks within cells to alter layout or alignment
% - do not nest tabular environments (no tabular environments within tabular environments)
% - no graphics or colored text (cell background color/shading OK)
% See http://journals.plos.org/plosone/s/tables for table guidelines.
%
% For tables that exceed the width of the text column, use the adjustwidth environment as illustrated in the example table in text below.
%
% % % % % % % % % % % % % % % % % % % % % % % %
%
% -- EQUATIONS, MATH SYMBOLS, SUBSCRIPTS, AND SUPERSCRIPTS
%
% IMPORTANT
% Below are a few tips to help format your equations and other special characters according to our specifications. For more tips to help reduce the possibility of formatting errors during conversion, please see our LaTeX guidelines at http://journals.plos.org/plosone/s/latex
%
% For inline equations, please be sure to include all portions of an equation in the math environment.  For example, x$^2$ is incorrect; this should be formatted as $x^2$ (or $\mathrm{x}^2$ if the romanized font is desired).
%
% Do not include text that is not math in the math environment. For example, CO2 should be written as CO\textsubscript{2} instead of CO$_2$.
%
% Please add line breaks to long display equations when possible in order to fit size of the column. 
%
% For inline equations, please do not include punctuation (commas, etc) within the math environment unless this is part of the equation.
%
% When adding superscript or subscripts outside of brackets/braces, please group using {}.  For example, change "[U(D,E,\gamma)]^2" to "{[U(D,E,\gamma)]}^2". 
%
% Do not use \cal for caligraphic font.  Instead, use \mathcal{}
%
% % % % % % % % % % % % % % % % % % % % % % % % 
%
% Please contact with any questions.
%
% % % % % % % % % % % % % % % % % % % % % % % %

\documentclass[10pt,letterpaper]{article}
\usepackage[top=0.85in,left=2.75in,footskip=0.75in]{geometry}

% amsmath and amssymb packages, useful for mathematical formulas and symbols
\usepackage{amsmath,amssymb}

% Use adjustwidth environment to exceed column width (see example table in text)
\usepackage{changepage}
\usepackage{tabularx}
\usepackage{array}
\usepackage{makecell}
\usepackage{adjustbox}
\usepackage{float}
\usepackage{booktabs}
\usepackage{dirtree}

% textcomp package and marvosym package for additional characters
\usepackage{textcomp,marvosym}

% cite package, to clean up citations in the main text. Do not remove.
\usepackage{cite}

% Use nameref to cite supporting information files (see Supporting Information section for more info)
\usepackage{nameref,hyperref}

% line numbers
\usepackage[right]{lineno}

% ligatures disabled
\usepackage[nopatch=eqnum]{microtype}
\DisableLigatures[f]{encoding = *, family = * }

% color can be used to apply background shading to table cells only
\usepackage[table]{xcolor}

% array package and thick rules for tables
\usepackage{array}

% create "+" rule type for thick vertical lines
\newcolumntype{+}{!{\vrule width 2pt}}

% create \thickcline for thick horizontal lines of variable length
\newlength\savedwidth
\newcommand\thickcline[1]{%
  \noalign{\global\savedwidth\arrayrulewidth\global\arrayrulewidth 2pt}%
  \cline{#1}%
  \noalign{\vskip\arrayrulewidth}%
  \noalign{\global\arrayrulewidth\savedwidth}%
}

% \thickhline command for thick horizontal lines that span the table
\newcommand\thickhline{\noalign{\global\savedwidth\arrayrulewidth\global\arrayrulewidth 2pt}%
\hline
\noalign{\global\arrayrulewidth\savedwidth}}


% Remove comment for double spacing
%\usepackage{setspace} 
%\doublespacing

% Text layout
\raggedright
\setlength{\parindent}{0.5cm}
\textwidth 5.25in
\textheight 8.75in

% Bold the 'Figure #' in the caption and separate it from the title/caption with a period
% Captions will be left justified
\usepackage[aboveskip=1pt,labelfont=bf,labelsep=period,justification=raggedright,singlelinecheck=off]{caption}
\renewcommand{\figurename}{Fig}

% Please use the included `plos2025.bst` as your BibTeX style
\bibliographystyle{plos2025}

% Remove brackets from numbering in List of References
\makeatletter
\renewcommand{\@biblabel}[1]{\quad#1.}
\makeatother


% Header and Footer with logo
\usepackage{lastpage,fancyhdr,graphicx}
\usepackage{epstopdf}
%\pagestyle{myheadings}
\pagestyle{fancy}
\fancyhf{}
%\setlength{\headheight}{27.023pt}
%\lhead{\includegraphics[width=2.0in]{PLOS-submission.eps}}
\rfoot{\thepage/\pageref{LastPage}}
\renewcommand{\headrulewidth}{0pt}
\renewcommand{\footrule}{\hrule height 2pt \vspace{2mm}}
\fancyheadoffset[L]{2.25in}
\fancyfootoffset[L]{2.25in}
\lfoot{\today}

%% Include all user-defined macros below

\newcommand{\lorem}{\textbf{LOREM}}
\newcommand{\ipsum}{\textbf{IPSUM}}

%% END MACROS SECTION


\begin{document}
\vspace*{0.2in}

% Title must be 250 characters or less.
\begin{flushleft}
{\Large
\textbf{Large-Scale Statistical Dissection of Sequence-Derived Biochemical Features Distinguishing Soluble and Insoluble Proteins} % Please use "sentence case" for title and headings (capitalize only the first word in a title (or heading), the first word in a subtitle (or subheading), and any proper nouns).
}
\newline
% Insert author names, affiliations and corresponding author email (do not include titles, positions, or degrees).
\\
Nguyen Huy Hoang Vu
\textsuperscript{1},
Bao Long Nguyen
\textsuperscript{2,*}
\\
\bigskip
\textbf{1} Faculty of Mathematics, Mechanics and Informatics, University of Science, Vietnam National University, Hanoi (VNU-HUS), Hanoi, Vietnam
\\
\textbf{2} School of Economics and Management, Hanoi University of Science and Technology (HUST), Hanoi, Vietnam
\\
\bigskip

% Insert additional author notes using the symbols described below. Insert symbol callouts after author names as necessary.
% 
% Remove or comment out the author notes below if they aren't used.
%
% Primary Equal Contribution Note

% Additional Equal Contribution Note
% Also use this double-dagger symbol for special authorship notes, such as senior authorship.

% Current address notes
% change symbol to "\textcurrency a" if more than one current address note
% \textcurrency b Insert second current address 
% \textcurrency c Insert third current address

% Deceased author note

% Group/Consortium Author Note

% Use the asterisk to denote corresponding authorship and provide email address in note below.
*

\end{flushleft}
% Please keep the abstract below 300 words

% For PLOS Medicine research article authors, please structure your abstract
% with "Background", "Method and Findings" and "Conclusion" sections per
% journal requirements.

% For PLOS Neglected Tropical Diseases research article authors, please
% structure your abstract with "Background", "Methodology", "Findings", and
% "Conclusion" sections per journal requirements.
%
\section*{Abstract}
Protein solubility critically influences recombinant expression efficiency and downstream biotechnological applications. While deep learning models have improved predictive accuracy, the intrinsic magnitude, redundancy, and interpretability of classical sequence-derived determinants remain insufficiently characterized. We performed a large-scale univariate analysis on a curated dataset of 78{,}031 proteins (46{,}450 soluble; 31{,}581 insoluble). Thirty-six biochemical descriptors were evaluated using Mann-Whitney U tests with Benjamini-Hochberg false discovery rate correction. Effect sizes were quantified using Cliff's $\delta$, and discriminative performance was assessed by ROC AUC. Although 34 features remained statistically significant after correction, most exhibited small effect sizes and substantial overlap between classes. The strongest effects were associated with size-related features (sequence length and molecular weight; $\delta \approx -0.21$), whereas charge-related descriptors, particularly the proportion of negatively charged residues ($\delta = 0.150$; AUC = 0.575), showed consistent but modest shifts. Spearman correlation analysis revealed near-complete redundancy among major size-related variables ($\rho$ up to 0.998). Applying a redundancy threshold ($|\rho| \ge 0.85$), we derived a parsimonious composite integrating sequence length and negative charge proportion, achieving AUC = 0.624 (MCC = 0.1746). These findings suggest that sequence-level solubility information is consistent with a low-dimensional organization at the level of global sequence-derived descriptors and governed by coordinated weak effects, establishing a transparent statistical baseline for large-scale solubility characterization.

% Please keep the Author Summary between 150 and 200 words. Use first person.
% PLOS ONE, PLOS Biology, PLOS Global Public Health, PLOS Mental Health, and PLOS Water authors please skip this step. Author Summary is not valid for submissions to these journals.

% For PLOS Medicine authors, please structure your author summary with answers to the following questions:
% Why was this study done?
% What did the researchers do and find?
% What do these findings mean?
%

\linenumbers

% Use "Eq" instead of "Equation" for equation citations.
\section*{Introduction}

Protein solubility is a fundamental physicochemical property that governs protein folding, intracellular stability, recombinant expression efficiency, and downstream biotechnological applications \cite{baneyx1999expression,chiti2017misfolding,rosano2014ecoli}. In heterologous expression systems, poor solubility frequently results in aggregation, inclusion body formation, reduced functional yield, and substantial experimental cost \cite{wilkinson1991solubility,chiti2006aggregation}. From a molecular perspective, solubility reflects the balance between intermolecular attractive forces that promote aggregation and intramolecular stabilizing interactions that favor properly folded conformations. Understanding how this balance is encoded in the primary amino acid sequence therefore remains a central problem in computational and structural biology.

A substantial body of experimental and computational work has shown that protein solubility is influenced by multiple sequence-derived properties. Chain length affects folding complexity, translational burden, and the likelihood of partially folded intermediates. Amino acid composition shapes the overall physicochemical landscape, including hydrophobicity, polarity, and charge distribution. Electrostatic interactions contribute to colloidal stability, while hydrophobic clustering and aggregation-prone segments facilitate intermolecular association \cite{idicula2005structure,ventura2005aggregation}. Importantly, these determinants are not independent: many classical descriptors are structurally or mathematically coupled, suggesting that observed effects may reflect shared latent physicochemical axes rather than distinct mechanisms.

Early computational approaches captured these determinants using interpretable sequence-derived descriptors such as amino acid composition, secondary structure propensity scales, and hydropathy indices \cite{chou1974prediction,kyte1982simple}. These features remain attractive due to their transparency and computational efficiency. However, their relative contribution, redundancy structure, and practical magnitude remain insufficiently characterized at large-scale. In high-powered datasets, extremely small $p$-values can arise from negligible distributional shifts, making it difficult to distinguish statistical significance from biological relevance \cite{wasserstein2019pvalue}.

Recent advances in machine-learning, particularly deep neural networks and protein language models, have achieved strong predictive performance in solubility benchmarks \cite{khurana2018deepsol,hon2021soluprot,thumuluri2022netsolp,zhang2024plmsol}. Despite their accuracy, these approaches often operate as high-capacity black boxes, obscuring the marginal contribution and interaction structure of individual physicochemical features. Moreover, such models typically require substantial computational resources, limiting their applicability in experimental settings where rapid and accessible prediction is needed.

A key unresolved question is whether sequence-level determinants of solubility act primarily through additive contributions of independent physicochemical factors, or whether they depend on higher-order sequence patterns that cannot be captured by classical descriptors. While higher-order representations may improve predictive performance, they introduce additional modeling layers, increase computational complexity, and may accumulate prediction error when inferred from primary sequence alone.

In parallel, many widely accepted determinants of solubility have been identified in relatively small or model-specific datasets, and their importance is often interpreted qualitatively. As a result, their true effect size, redundancy, and practical discriminative contribution remain unclear at scale. This motivates a systematic and statistically rigorous re-evaluation of classical sequence-derived features in a large and heterogeneous dataset.

In the present study, we perform a large-scale statistical dissection of 36 sequence-derived biochemical descriptors using a curated benchmark of soluble and insoluble proteins. Rather than developing a high-capacity predictive model, our objective is to quantify effect magnitude, redundancy structure, and practical discrimination under strict statistical control. By combining non-parametric testing, FDR correction, effect size estimation, ROC analysis, and correlation-based redundancy analysis, we examine whether protein solubility is driven by dominant single determinants or by coordinated weak signals across a low-dimensional physicochemical space.

This study provides a statistically principled and interpretable baseline framework for protein solubility derived from primary sequence features. By integrating large-scale effect size characterization with a reproducible analytical pipeline, it quantifies the practical magnitude and limitations of classical biochemical descriptors. Rather than proposing a higher-capacity predictor, the framework helps characterize the level of discrimination achievable within the restricted space of global sequence-derived physicochemical features, providing a transparent reference for assessing the added value of more complex predictive models.

Importantly, this study is not intended to develop or benchmark a predictive model. Instead, the objective is to quantify the intrinsic statistical structure of sequence-derived physicochemical features under minimal modeling assumptions.
Any discriminative performance reported in this work should therefore be interpreted as a descriptive property of the feature space rather than as an estimate of predictive generalization.

\section*{Materials and methods}
\subsection*{Dataset}

The dataset was obtained from the curated solubility benchmark introduced by Zhang \textit{et al.} (2024) \cite{zhang2024plmsol}, comprising more than 78{,}000 protein sequences annotated as soluble or insoluble. The original release provides independent training, validation, and test splits constructed for predictive benchmarking, distributed as three FASTA files containing approximately 74{,}000, 2{,}000, and 2{,}000 protein sequences, respectively.

For the present analysis, all splits were merged into a single dataset because the objective was large-scale distributional characterization rather than predictive generalization. The final dataset comprised 78{,}031 protein sequences (46{,}450 soluble and 31{,}581 insoluble). All sequences and labels are publicly available via Zenodo \cite{zhang2024plmsol}. Class imbalance in the dataset is unlikely to materially affect rank-based statistics such as the Mann–Whitney U test or ROC AUC.

Protein sequences are represented as contiguous strings of single-letter amino acid codes (e.g., ``ACDEFGHIK...'') without gaps or alignment. No additional filtering or preprocessing was applied, as the dataset had already been curated in the original benchmark study. In particular, no filtering based on sequence length, redundancy, or sequence identity was performed, and all sequences provided in the original dataset were retained.

Two sequences containing rare or ambiguous amino acid symbols (``X'') were preserved. These residues were excluded from amino acid frequency calculations but did not affect other sequence-derived descriptors. All features were computed directly from the raw FASTA sequences.

The FASTA files used in this study correspond exactly to those provided in the original benchmark without modification.

\subsection*{Feature Extraction}

All protein sequences were retained in their original form; ambiguous residues were preserved in the raw sequences but excluded from relevant feature calculations as described below. Feature extraction was performed using the 20 canonical amino acids; rare or ambiguous residues (e.g., ``X'') were not removed from the sequences but were excluded from frequency-based calculations to avoid distortion of compositional features. The presence of such residues was minimal and did not affect global descriptors beyond amino acid frequency measures.

A total of 36 sequence-derived biochemical features were computed for each protein, including:

\begin{itemize}
    \item \textbf{Amino acid frequency features (20 variables).}  
    For each amino acid $a \in \mathcal{A}$ (20 canonical residues), the frequency is defined as:
    \[
    f_a = \frac{n_a}{L},
    \]
    where $n_a$ is the number of occurrences of residue $a$ in the sequence and $L$ is the sequence length (excluding ambiguous residues such as ``X''). These variables quantify compositional bias at the residue level.

    \item \textbf{Functional residue group ratios.}  
    Residues were grouped according to physicochemical properties, including:
    positively charged ($\{K,R,H\}$), negatively charged ($\{D,E\}$), polar ($\{S,T,N,Q\}$), 
    hydrophobic ($\{A,V,I,L,M,F,W,Y\}$), small ($\{A,G,S,T\}$), and sulfur-containing ($\{C,M\}$).  
    For each group $G$, the ratio is defined as:
    \[
    r_G = \frac{\sum_{a \in G} n_a}{L}.
    \]
    These descriptors capture coarse-grained physicochemical composition.

    \item \textbf{Global physicochemical descriptors.}  
    Molecular weight was computed as the sum of residue masses.  
    The isoelectric point (pI) and net charge at pH 7 were estimated using standard residue pKa values and the Henderson–Hasselbalch equation.  
    Mean hydropathy was computed as the average Kyte–Doolittle hydropathy index across the sequence \cite{kyte1982simple}. These variables characterize global physicochemical properties of the sequence.

    \item \textbf{Secondary structure propensity proxies.}  
    Helix, sheet, and turn propensities were approximated by averaging residue-level Chou–Fasman parameters across the sequence \cite{chou1974prediction}.  
    For a given structural class $s$, the feature is:
    \[
    P_s = \frac{1}{L} \sum_{i=1}^{L} p_s(a_i),
    \]
    where $p_s(a_i)$ is the Chou–Fasman propensity of residue $a_i$ for structure $s$.

    \item \textbf{Intrinsic disorder proxy.}  
    Intrinsic disorder tendency was approximated using residue-level disorder propensity scales derived from established literature \cite{dunker2001disorder,uversky2013intrinsic}.  
    The disorder ratio was computed as the fraction of residues exceeding a predefined disorder propensity threshold:
    \[
    r_{\text{disorder}} = \frac{1}{L} \sum_{i=1}^{L} \mathbf{1}(d(a_i) > \tau),
    \]
    where $d(a_i)$ denotes the disorder score and $\tau$ is a fixed threshold. The exact propensity scale and threshold used are provided in Table S1 and in the accompanying code repository.

    \item \textbf{Aggregation-related proxy.}  
    Aggregation propensity was approximated by the longest contiguous hydrophobic segment in the sequence.  
    Let $\ell_{\max}$ denote the length of the longest run of residues in the hydrophobic set. The feature is defined as:
    \[
    r_{\text{agg}} = \frac{\ell_{\max}}{L}.
    \]
    This proxy captures local clustering of hydrophobic residues associated with aggregation risk.
\end{itemize}

These descriptors form a structured feature matrix for subsequent statistical evaluation.

\subsection*{Analytical workflow}

The analytical workflow was designed to quantify practical effect magnitude, control false discoveries, eliminate redundancy, and subsequently construct an interpretable composite index.

For each of the 36 sequence-derived descriptors, distributional differences between soluble and insoluble proteins were first evaluated using the Mann-Whitney U test \cite{mann1947test}. Resulting $p$-values were adjusted via the Benjamini-Hochberg procedure to control the false discovery rate (FDR) \cite{benjamini1995controlling}. Effect sizes (Cliff’s $\delta$), which measure stochastic dominance without distributional assumptions \cite{cliff1993dominance}, were computed for all features. For features meeting FDR significance, effect magnitudes were interpreted alongside confidence intervals. Median shifts were expressed using the Hodges–Lehmann estimator with 95\% confidence intervals \cite{hodges1963estimates}. Stability of $\delta$ was assessed via percentile bootstrap resampling ($B = 2{,}000$) \cite{efron1993bootstrap}. Univariate discriminative capacity was evaluated using ROC AUC \cite{hanley1982roc}, with optimal thresholds determined by Youden’s $J$ statistic \cite{youden1950index}.

Features were subsequently ranked according to absolute $\delta$ to identify descriptors exhibiting the strongest practical separation. Because high-effect features may capture overlapping physicochemical dimensions, pairwise redundancy was assessed prior to composite modeling using Spearman’s rank correlation coefficient \cite{spearman1904,conover1999}. Spearman correlation was selected due to its robustness to non-normality and its ability to capture monotonic relationships.

Feature pairs exhibiting strong monotonic association ($|\rho| \ge 0.85$) were considered redundant. Within correlated clusters, only a single representative descriptor was retained to avoid double-counting latent physicochemical axes. This redundancy-aware filtering ensured that subsequent integration combined orthogonal dimensions rather than correlated proxies of the same underlying property.

The final composite-$\delta$ index was constructed as a linear combination of retained features, robustly scaled using the median and interquartile range (IQR) \cite{Huber1981,Hampel1986}, and weighted by their corresponding $\delta$ values. Proteins were classified according to the sign of the composite score.

The composite-$\delta$ index is not a trained predictive model. Its construction is fully data-dependent and based on empirical effect size estimates derived from the same dataset. Because no parameter fitting or hyperparameter optimization is performed, the resulting index summarizes distributional separability under a statistically controlled and redundancy-filtered feature set.

Consequently, any performance metrics (e.g., AUC, MCC) reported for this index do not represent out-of-sample predictive performance, but rather quantify descriptive separability within the observed feature space.

\begin{figure}[H]
\centering
\includegraphics[width=0.4\linewidth]{Analytical workflow.png}
\caption{Analytical workflow.}
\label{fig:pipeline_en}
\end{figure}

% Results and Discussion can be combined.
\section*{Results}
\subsection*{Dataset overview and multiple-testing adjusted significance}

Following preprocessing and feature extraction, the final analysis dataset comprised $N = 78{,}031$ protein sequences, including $46{,}450$ soluble and $31{,}581$ insoluble entries. For each sequence, 36 sequence-derived biochemical descriptors were computed, spanning composition, charge-related variables, hydrophobicity measures, structural propensities, and aggregation proxies.

Group-wise distributional differences were evaluated using the Mann-Whitney U test \cite{mann1947test}, with Benjamini-Hochberg correction to control the false discovery rate (FDR) \cite{benjamini1995controlling}. After adjustment, $34$ of $36$ features remained statistically significant ($q < 0.05$), indicating that most classical sequence-level descriptors exhibit detectable distributional shifts between soluble and insoluble proteins.

However, given the large sample size, statistical significance alone is insufficient to infer biological relevance. In high-powered datasets, even negligible shifts can produce extremely small $p$-values \cite{wasserstein2019pvalue}. We therefore focus on two complementary quantities: (i) effect magnitude quantified by Cliff’s $\delta$ \cite{cliff1993dominance}, which measures stochastic dominance independent of distributional assumptions, and (ii) univariate discriminative capacity assessed via ROC AUC \cite{hanley1982roc}, which provides a descriptive measure of class separability.

To express group differences on the original measurement scale, we report the Hodges–Lehmann (HL) location shift with 95\% confidence intervals \cite{hodges1963estimates}. Uncertainty of $\delta$ estimates was evaluated using percentile bootstrap intervals ($B = 2{,}000$) \cite{efron1993bootstrap}. For completeness, Youden’s $J$ statistic and its corresponding optimal threshold $T$ are also reported \cite{youden1950index}; in the presence of substantial distributional overlap, $J$ remains close to zero even when statistical significance is achieved.

Collectively, this framework distinguishes statistical detectability from practical magnitude, enabling a biologically grounded interpretation of whether sequence-derived physicochemical features exert strong or merely subtle influence on protein solubility.

\subsection*{Global physicochemical features}

\begin{table}[H]
\centering
\small
\caption{Results for 16 global sequence-derived features.}
\setlength{\tabcolsep}{3pt}
\renewcommand{\arraystretch}{1.15}
\begin{adjustbox}{max width=\textwidth}
\begin{tabular}{lccccccll}
\toprule
\textbf{Feature} &
\textbf{Soluble (median [Q1,Q3])} &
\textbf{Insoluble (median [Q1,Q3])} &
\textbf{HL [95\% CI]} &
$\boldsymbol{\delta}$ \textbf{[95\% CI]} &
\textbf{FDR $q$} &
\textbf{AUC} &
\textbf{$J$} &
\textbf{$T^\ast$} \\
\midrule

length
& 209 [129, 338]
& 277 [172, 426]
& -70 [-77, -62]
& -0.215 [-0.237, -0.196]
& 0.000
& 0.392
& 0.158
& 227 \\

molecular\_weight
& 19\,743 [12\,201, 31\,715]
& 25\,966 [16\,148, 39\,869]
& -6322.485 [-7147.547, -5784.646]
& -0.214 [-0.244, -0.204]
& 0.000
& 0.393
& 0.157
& 21520 \\

aggregation\_ratio
& 0.02262 [0.01502, 0.03371]
& 0.01869 [0.01252, 0.02857]
& 0.004 [0.003, 0.004]
& 0.166 [0.138, 0.178]
& 0.000
& 0.583
& 0.125
& 0.020 \\

neg\_ratio
& 0.1298 [0.1104, 0.1510]
& 0.1219 [0.1034, 0.1414]
& 0.008 [0.007, 0.009]
& 0.150 [0.134, 0.173]
& 0.000
& 0.575
& 0.114
& 0.126 \\

isoelectric\_point
& 6.401 [4.971, 8.357]
& 6.931 [5.384, 8.941]
& -0.588 [-0.702, -0.468]
& -0.132 [-0.167, -0.125]
& 0.000
& 0.434
& 0.102
& 6.523 \\

net\_charge\_pH7
& -1.882 [-7.066, 3.770]
& -0.207 [-5.000, 6.476]
& -1.618 [-1.931, -1.217]
& -0.120 [-0.143, -0.099]
& 0.000
& 0.440
& 0.107
& 2.413 \\

mean\_hydropathy
& -0.3511 [-0.4695, -0.2277]
& -0.3085 [-0.4388, -0.1834]
& -0.046 [-0.060, -0.035]
& -0.080 [-0.106, -0.064]
& 0.000
& 0.460
& 0.067
& -0.305 \\

sulfur\_ratio
& 0.03448 [0.01961, 0.04712]
& 0.03681 [0.02193, 0.05013]
& -0.002 [-0.003, -0.002]
& -0.076 [-0.096, -0.056]
& 0.000
& 0.462
& 0.066
& 0.029 \\

helix\_prop\_mean
& 1.029 [1.019, 1.038]
& 1.025 [1.015, 1.034]
& 0.004 [0.003, 0.005]
& 0.060 [0.049, 0.089]
& 0.000
& 0.530
& 0.054
& 1.041 \\

hydrophobic\_ratio
& 0.4019 [0.3786, 0.4253]
& 0.4063 [0.3829, 0.4286]
& -0.003 [-0.005, -0.001]
& -0.051 [-0.053, -0.012]
& 0.000
& 0.475
& 0.046
& 0.419 \\

sheet\_prop\_mean
& 0.9937 [0.9857, 1.0024]
& 0.9963 [0.9883, 1.0048]
& -0.002 [-0.003, -0.000]
& -0.050 [-0.056, -0.015]
& 0.000
& 0.475
& 0.046
& 1.008 \\

disorder\_ratio
& 0.3976 [0.3690, 0.4310]
& 0.3937 [0.3662, 0.4274]
& 0.003 [0.001, 0.005]
& 0.048 [0.021, 0.063]
& 0.000
& 0.524
& 0.039
& 0.412 \\

polar\_ratio
& 0.2006 [0.1765, 0.2247]
& 0.2031 [0.1798, 0.2278]
& -0.002 [-0.005, -0.001]
& -0.024 [-0.055, -0.013]
& 0.000
& 0.488
& 0.023
& 0.195 \\

tiny\_ratio
& 0.1389 [0.1111, 0.1667]
& 0.1411 [0.1127, 0.1695]
& -0.000 [-0.003, 0.002]
& -0.021 [-0.035, 0.009]
& 0.000
& 0.489
& 0.037
& 0.171 \\

pos\_ratio
& 0.1412 [0.1212, 0.1636]
& 0.1419 [0.1230, 0.1636]
& -0.000 [-0.002, 0.001]
& -0.015 [-0.026, 0.015]
& 0.000
& 0.492
& 0.022
& 0.116 \\

turn\_prop\_mean
& 0.9667 [0.9584, 0.9759]
& 0.9672 [0.9589, 0.9763]
& -0.001 [-0.003, 0.000]
& -0.011 [-0.036, 0.004]
& 0.011
& 0.495
& 0.021
& 0.928 \\

\bottomrule
\end{tabular}
\end{adjustbox}
\label{tab:global_features}
\end{table}

Table~\ref{tab:global_features} summarizes 16 global and grouped-composition descriptors. The largest absolute effects were observed for \texttt{length} and \texttt{molecular\_weight}, both exhibiting $\delta < 0$, indicating that insoluble proteins are, on average, longer and heavier. On the original measurement scale, the Hodges–Lehmann estimates correspond to a median shift of $-70$ amino acids (95\% CI $[-77,-62]$) for \texttt{length} and $-6{,}322$ Da (95\% CI $[-7{,}148,-5{,}785]$) for \texttt{molecular\_weight}. 

From a mechanistic standpoint, increased chain length elevates folding complexity, prolongs exposure of partially folded intermediates, and increases the probability of intermolecular encounters that nucleate aggregation \cite{baneyx1999expression,wilkinson1991solubility}. Larger proteins also present greater solvent-accessible surface area and a higher combinatorial risk of hydrophobic clustering. Despite these biologically plausible trends, univariate discrimination remained limited ($\mathrm{AUC} = 0.392$ and $0.393$, respectively). Values below 0.5 reflect reversed directionality rather than meaningful separability; indeed, inverting the decision rule yields $\mathrm{AUC} \approx 0.607$, yet substantial distributional overlap persists, as confirmed by small Youden’s $J$ statistics.

Charge-related descriptors exhibited coherent and biologically interpretable shifts. The proportion of negatively charged residues (\texttt{neg\_ratio}) was enriched in soluble proteins ($\delta = 0.150$, 95\% CI $[0.134,0.173]$, $\mathrm{AUC} = 0.575$; HL $=0.00810$ with 95\% CI $[0.00674,0.00937]$). Conversely, \texttt{isoelectric\_point} and \texttt{net\_charge\_pH7} showed $\delta < 0$, indicating higher $pI$ and reduced net negativity among insoluble proteins. These findings align with electrostatic stabilization theory: increased net negative charge enhances intermolecular repulsion, reduces colloidal attraction, and suppresses aggregation propensity \cite{wilkinson1991solubility,idicula2005structure,chiti2006aggregation}. The moderate AUC values ($\approx 0.57$) indicate measurable but limited standalone discriminatory power.

Hydrophobicity-related descriptors exhibited smaller effects. Both \texttt{mean\_hydropathy} and \texttt{hydrophobic\_ratio} showed $\delta < 0$, consistent with the role of exposed hydrophobic patches in promoting solvent exclusion and intermolecular association \cite{kyte1982simple,ventura2005aggregation}. However, effect sizes were modest and AUC values remained close to 0.5, underscoring extensive overlap between soluble and insoluble distributions. These observations suggest that global hydrophobic averages are insufficient to capture context-dependent aggregation mechanisms.

\subsection*{Disorder and secondary structure proxies}

Intrinsic disorder tendency, approximated by \texttt{disorder\_ratio}, exhibited a small positive effect ($\delta = 0.048$, $\mathrm{AUC} = 0.524$), suggesting slight enrichment of disorder-associated residues among soluble proteins. Disordered or flexible regions may increase solvent accessibility and modulate intermolecular interaction landscapes, although their impact appears context-dependent and modest in magnitude \cite{dunker2001disorder,uversky2013intrinsic}. 

Secondary structure propensity proxies (\texttt{helix\_prop\_mean}, \texttt{sheet\_prop\_mean}, and \texttt{turn\_prop\_mean}) demonstrated very small effect sizes. This indicates that simple global aggregation of Chou-Fasman propensities across full-length sequences does not yield strong univariate separation in a large and heterogeneous dataset \cite{chou1974prediction}.

\subsection*{Amino acid composition}

\begin{table}[H]
\centering
\small
\caption{Results for amino acid frequency features.}
\setlength{\tabcolsep}{3pt}
\renewcommand{\arraystretch}{1.15}
\begin{adjustbox}{max width=\textwidth}
\begin{tabular}{lccccccll}
\toprule
\textbf{Feature} &
\textbf{Soluble (median [Q1,Q3])} &
\textbf{Insoluble (median [Q1,Q3])} &
\textbf{HL [95\% CI]} &
$\boldsymbol{\delta}$ \textbf{[95\% CI]} &
\textbf{FDR $q$} &
\textbf{AUC} &
\textbf{$J$} &
\textbf{$T^\ast$} \\
\midrule

freq\_C & 0.01017 [0.00312, 0.01882] & 0.01282 [0.00615, 0.02143]
& -0.003 [-0.003, -0.002]
& -0.131 [-0.151, -0.111]
& 0.000
& 0.434
& 0.100
& 0.000 \\

freq\_E & 0.07064 [0.05488, 0.08889] & 0.06529 [0.05072, 0.08187]
& 0.006 [0.005, 0.007]
& 0.121 [0.119, 0.160]
& 0.000
& 0.560
& 0.092
& 0.071 \\

freq\_R & 0.05350 [0.03663, 0.07273] & 0.05894 [0.04061, 0.07944]
& -0.005 [-0.006, -0.005]
& -0.105 [-0.101, -0.061]
& 0.000
& 0.447
& 0.083
& 0.060 \\

freq\_K & 0.06019 [0.04255, 0.07843] & 0.05350 [0.03636, 0.07339]
& 0.007 [0.006, 0.008]
& 0.091 [0.067, 0.109]
& 0.000
& 0.546
& 0.080
& 0.049 \\

freq\_D & 0.05721 [0.04098, 0.07391] & 0.05473 [0.04000, 0.07074]
& 0.003 [0.002, 0.003]
& 0.080 [0.055, 0.097]
& 0.000
& 0.540
& 0.067
& 0.061 \\

freq\_S & 0.06349 [0.04724, 0.08065] & 0.06630 [0.05000, 0.08333]
& -0.003 [-0.003, -0.002]
& -0.068 [-0.092, -0.044]
& 0.000
& 0.466
& 0.043
& 0.067 \\

freq\_P & 0.04244 [0.02899, 0.05755] & 0.04494 [0.03125, 0.06061]
& -0.003 [-0.003, -0.002]
& -0.064 [-0.088, -0.040]
& 0.000
& 0.468
& 0.050
& 0.048 \\

freq\_A & 0.07273 [0.05556, 0.09091] & 0.07650 [0.05825, 0.09524]
& -0.004 [-0.005, -0.003]
& -0.051 [-0.075, -0.026]
& 0.000
& 0.474
& 0.052
& 0.092 \\

freq\_Q & 0.03768 [0.02500, 0.05172] & 0.03643 [0.02424, 0.05000]
& 0.001 [0.001, 0.002]
& 0.042 [0.017, 0.066]
& 0.000
& 0.521
& 0.042
& 0.030 \\

freq\_Y & 0.03061 [0.01923, 0.04211] & 0.02913 [0.01818, 0.04110]
& 0.001 [0.001, 0.002]
& 0.041 [0.016, 0.066]
& 0.000
& 0.521
& 0.041
& 0.009 \\

freq\_W & 0.01000 [0.00000, 0.01754] & 0.01031 [0.00000, 0.01852]
& -0.000 [-0.001, 0.000]
& -0.039 [-0.063, -0.014]
& 0.000
& 0.481
& 0.047
& 0.000 \\

freq\_I & 0.05128 [0.03604, 0.06818] & 0.05333 [0.03788, 0.07059]
& -0.002 [-0.003, -0.001]
& -0.039 [-0.063, -0.014]
& 0.000
& 0.481
& 0.047
& 0.045 \\

freq\_V & 0.06140 [0.04545, 0.07813] & 0.06349 [0.04688, 0.08000]
& -0.002 [-0.003, -0.001]
& -0.035 [-0.060, -0.010]
& 0.000
& 0.482
& 0.043
& 0.077 \\

freq\_N & 0.03922 [0.02632, 0.05263] & 0.04000 [0.02740, 0.05405]
& -0.001 [-0.002, -0.000]
& -0.033 [-0.058, -0.008]
& 0.000
& 0.484
& 0.040
& 0.056 \\

freq\_G & 0.06195 [0.04598, 0.07937] & 0.06383 [0.04688, 0.08108]
& -0.002 [-0.003, -0.001]
& -0.028 [-0.053, -0.003]
& 0.000
& 0.487
& 0.039
& 0.031 \\

freq\_L & 0.09091 [0.07317, 0.10976] & 0.09195 [0.07407, 0.11111]
& -0.001 [-0.002, -0.000]
& -0.026 [-0.051, -0.001]
& 0.000
& 0.487
& 0.038
& 0.079 \\

freq\_F & 0.03636 [0.02532, 0.04878] & 0.03704 [0.02632, 0.05000]
& -0.001 [-0.001, -0.000]
& -0.018 [-0.043, 0.007]
& 0.000
& 0.491
& 0.029
& 0.018 \\

freq\_H & 0.02381 [0.01389, 0.03448] & 0.02439 [0.01471, 0.03571]
& -0.001 [-0.001, 0.000]
& -0.016 [-0.041, 0.009]
& 0.000
& 0.492
& 0.029
& 0.012 \\

freq\_M & 0.02174 [0.01220, 0.03125] & 0.02174 [0.01282, 0.03125]
& 0 [-0.001, 0.001]
& -0.003 [-0.028, 0.023]
& 0.514
& 0.499
& 0.023
& 0.011 \\

freq\_T & 0.05263 [0.03704, 0.06897] & 0.05263 [0.03788, 0.06897]
& 0 [-0.001, 0.001]
& 0.000 [-0.025, 0.025]
& 0.989
& 0.500
& 0.022
& 0.072 \\

\bottomrule
\end{tabular}
\end{adjustbox}
\label{tab:aa_freq}
\end{table}
Table~\ref{tab:aa_freq} reports results for the 20 amino acid frequency variables. Many compositional descriptors achieved extremely small $q$-values, reflecting high statistical power; however, most displayed small $|\delta|$ and AUC values only marginally deviating from 0.5. Notably, \texttt{freq\_E} and \texttt{freq\_D} were elevated in soluble proteins, consistent with the \texttt{neg\_ratio} signal, whereas \texttt{freq\_R} and \texttt{freq\_C} were relatively enriched in insoluble proteins. These trends are compatible with electrostatic contributions and residue-specific side chain chemistry influencing folding stability and aggregation kinetics \cite{wilkinson1991solubility,idicula2005structure,ventura2005aggregation}. Only two features, \texttt{freq\_M} and \texttt{freq\_T}, did not remain significant after FDR correction ($q \ge 0.05$), with $\delta \approx 0$ and $\mathrm{AUC} \approx 0.5$, indicating negligible univariate discriminatory information within this dataset.

\subsection*{Emergent weak-signal regime and biological interpretation}

Taken together, these findings indicate that protein solubility is not governed by a dominant sequence-level determinant but instead reflects coordinated contributions of multiple weak physicochemical signals \cite{chiti2006aggregation,chiti2017misfolding}. Size-related variables capture a structural burden axis, while charge-related descriptors reflect electrostatic stabilization; hydrophobic and compositional features contribute smaller contextual effects. The generally small Youden’s $J$ values across individual descriptors confirm that no single feature provides practical threshold-based separation. Rather, the data support a weak-signal, potentially low-dimensional structure in which overlapping physicochemical axes jointly influence solubility. This observation motivates subsequent redundancy-aware integration of selected descriptors into a parsimonious composite index, as detailed below.

\begin{figure}[H]
\centering
\includegraphics[width=0.5\textwidth]{roc_top4.png}
\caption{Univariate ROC curves for the four features with the largest absolute Cliff’s $\delta$.}
\label{fig:roc}
\end{figure}

The observed asymmetry between negatively and positively charged residues warrants further interpretation. Enrichment of negatively charged residues in soluble proteins is consistent with established biophysical principles, as increased net negative charge enhances electrostatic repulsion and reduces intermolecular association, thereby promoting solubility \cite{kramer2012charge,chan2013solubility}.

In contrast, positively charged residues may participate in non-specific interactions with nucleic acids, membranes, or other cellular components, which can facilitate aggregation under certain conditions. 

It is important to note that the dataset is derived from recombinant expression in Escherichia coli, and therefore the observed patterns may partially reflect system-specific constraints. The intracellular environment and expression constraints of this system may introduce biases that favor negatively charged, more soluble constructs. Therefore, the observed effect likely reflects a combination of general physicochemical principles and system-specific expression biases. Generalization to other expression systems or native proteomes warrants further investigation.

\subsection*{Redundancy structure and composite-$\delta$ refinement}

\subsubsection*{Spearman redundancy analysis}

Prior to finalizing the composite formulation, pairwise redundancy among selected high-effect features was evaluated using Spearman’s rank correlation coefficient \cite{spearman1904}. The resulting correlation matrix is shown in Figure~\ref{fig:corr_matrix}.

\begin{figure}[H]
\centering
\includegraphics[width=0.6\linewidth]{corr_4features_spearman.png}
\caption{Spearman correlation matrix among top ranked global features.}
\label{fig:corr_matrix}
\end{figure}

Strong monotonic associations were observed among size-related descriptors. In particular, sequence length and molecular weight exhibited near-complete collinearity ($\rho \approx 0.998$), reflecting their deterministic structural coupling. Aggregation-related metrics also displayed strong monotonic association with size-related variables ($|\rho| \approx 0.9$), indicating that these descriptors largely capture a shared latent structural axis. 

In contrast, the proportion of negatively charged residues (\texttt{neg\_ratio}) showed minimal correlation with size-related variables ($|\rho| < 0.05$), suggesting relative independence of electrostatic and size dimensions. Applying a predefined redundancy criterion ($|\rho| \ge 0.85$), correlated size-related descriptors were considered redundant. To avoid double-counting a single latent physicochemical axis, only one representative size descriptor was retained. We selected \texttt{length} due to its direct structural interpretability and deterministic relation to molecular mass. The threshold ($|\rho| \ge 0.85$) was adopted as a conservative criterion to remove strongly collinear features while retaining distinct physicochemical dimensions. A formal dimensionality analysis (e.g., PCA) was not performed and is left for future work.

\begin{figure}[H]
\centering
\includegraphics[width=0.48\textwidth]{distribution length.png}
\includegraphics[width=0.48\textwidth]{distribution negative.png}
\caption{
Distributions of sequence length (left) and negative charge proportion (right) for soluble and insoluble proteins, showing substantial overlap between classes.
}
\label{fig:distribution}
\end{figure}

To illustrate the extent of distributional overlap, Figure~\ref{fig:distribution} presents the density distributions of the two principal orthogonal features: sequence length and negative charge proportion. In both cases, soluble and insoluble proteins exhibit substantial overlap despite statistically significant differences in central tendency.

For sequence length, insoluble proteins display a right-shifted distribution, reflecting a tendency toward longer chains. However, a considerable fraction of soluble proteins occupies the same range, limiting practical separability. Similarly, for negative charge proportion, soluble proteins show a modest enrichment in negatively charged residues, yet the two classes remain strongly intermixed.

These observations provide a direct visual confirmation that, although detectable at scale, the underlying physicochemical differences are small relative to within class variability. This substantial overlap explains the limited univariate discriminative performance observed in ROC analysis and reinforces the interpretation that protein solubility is governed by coordinated weak effects rather than strongly separable individual features. 

The initial composite-$\delta$ index integrated the top ranked descriptors by absolute effect size, robustly scaled using median and interquartile range (IQR) to mitigate the influence of outliers.

\begin{equation}
x_j^{(\mathrm{rs})}
=
\frac{x_j - \operatorname{median}(x_j)}
{\operatorname{IQR}(x_j)},
\qquad
\operatorname{IQR}(x_j)=Q_{0.75}(x_j)-Q_{0.25}(x_j)
\end{equation}

\begin{equation}
S
=
\sum_{j=1}^{k} \delta_j \, x_j^{(\mathrm{rs})}
\end{equation}

\begin{equation}
\hat{y} =
\begin{cases}
1, & \text{if } S > 0 \quad (\text{soluble}) \\
0, & \text{if } S \le 0 \quad (\text{insoluble})
\end{cases}
\end{equation}

\subsubsection*{Reduced composite formulation}

\begin{table}[H]
\centering
\caption{Robust summaries of the two retained physicochemical dimensions defining the reduced composite index}
\begin{tabular}{lc}
\hline
\textbf{Feature} & \textbf{Median [IQR]} \\
\hline
length 
& 236 [231] \\

neg\_ratio 
& 0.126531 [0.039934] \\

\hline
\end{tabular}
\label{tab:global_summary}
\end{table}

Using global medians and IQR values (Table~\ref{tab:global_summary}), the reduced composite-$\delta$ score becomes:

\begin{equation}
S =
-0.215\,\frac{L-236}{231}
+0.150\,\frac{N_{-}-0.126531}{0.039934}
\end{equation}

This two-dimensional formulation preserves the dominant size and electrostatic axes while eliminating redundant contributions. As shown below, discriminative performance remains comparable to the full multi-feature composite, supporting the interpretation that sequence-level solubility information is consistent with a low-dimensional organization at the level of global sequence-derived descriptors. The use of a linear formulation is intentional, as the objective is to isolate the contribution of first-order physicochemical features without introducing higher-order interactions that may obscure interpretability.

\begin{table}[H]
\centering
\small
\setlength{\tabcolsep}{3pt}
\renewcommand{\arraystretch}{1.15}
\caption{Contextual performance reference for existing solubility prediction methods and the composite-$\delta$ index.}
\begin{adjustbox}{max width=\textwidth}
\begin{tabular}{lcccccccc}
\toprule
\textbf{Model} & \textbf{$T$} & \textbf{AUC} & \textbf{Acc.} & \textbf{F1} & \textbf{MCC} & \textbf{Prec.} & \textbf{Sens.} & \textbf{Spec.} \\
\midrule
Protein\_sol & 0.5 & 0.5985 & 0.5649 & 0.6350 & 0.1405 & 0.5470 & 0.7569 & 0.3729 \\
SKADE & 0.5 & 0.6882 & 0.6443 & 0.5304 & 0.3305 & 0.7811 & 0.4015 & 0.8874 \\
SWI & 0.5 & 0.5597 & 0.5423 & 0.6334 & 0.0971 & 0.5284 & 0.7905 & 0.2938 \\
Solupro & 0.5 & 0.7126 & 0.6308 & 0.6639 & 0.2667 & 0.6095 & 0.7290 & 0.5325 \\
EPSOL & 0.5 & 0.6664 & 0.6663 & 0.5915 & 0.3578 & 0.7630 & 0.4830 & 0.8499 \\
NetSolP & 0.5 & 0.6183 & 0.5502 & 0.6557 & 0.1268 & 0.5312 & 0.8564 & 0.2437 \\
DeepSol-E & 0.4 & 0.6660 & 0.5990 & 0.6407 & 0.2036 & 0.5804 & 0.7150 & 0.4830 \\
PLM\_Sol & 0.5 & 0.8342 & 0.7299 & 0.7542 & 0.4690 & 0.6919 & 0.8289 & 0.6308 \\
\textbf{Composite-$\delta$} & 0 & \textbf{0.6240} & \textbf{0.5734} & \textbf{0.5875} & \textbf{0.1746} & \textbf{0.6922} & \textbf{0.5102} & \textbf{0.6663} \\
\bottomrule
\end{tabular}
\end{adjustbox}
\label{tab:model_comparison}
\end{table}

Performance metrics (AUC and MCC) of reference models were taken directly from the original benchmark study \cite{zhang2024plmsol} to provide contextual reference. These values are shown alongside the descriptive metrics of the composite-$\delta$ index but do not constitute a direct head-to-head comparison.

The comparison in Table \ref{tab:model_comparison} is provided for contextual reference only. The reported performance of the composite-$\delta$ index is not directly comparable to supervised models evaluated under strict train/test separation, as the present analysis does not involve independent validation. Instead, these values should be interpreted as descriptive indicators of signal strength in sequence-derived features.

Because the present study merges the original training, validation, and test splits for distributional analysis rather than predictive evaluation, the reported metrics should not be interpreted as estimates of out-of-sample performance.

Importantly, the proposed composite-$\delta$ index does not constitute a trained predictive model. Its construction is fully data-dependent and based on empirical effect size estimates derived from the same dataset. Consequently, AUC and MCC are reported solely as descriptive measures of separability within the observed feature space, rather than as indicators of predictive generalization.

The purpose of the composite-$\delta$ formulation is therefore not to achieve state-of-the-art predictive performance, but to provide an interpretable statistical reference that characterizes the magnitude and structure of sequence-derived solubility signals.

\begin{table}[H]
\centering
\small
\setlength{\tabcolsep}{3pt}
\renewcommand{\arraystretch}{1.15}
\caption{Qualitative comparison of inference-time computational requirements across representative solubility prediction approaches}
\begin{adjustbox}{max width=\textwidth}
\begin{tabular}{lccc}
\toprule
\textbf{Model} & \textbf{Architecture Type} & \textbf{Inference Complexity} & \textbf{Training Requirement} \\
\midrule
Protein\_sol & Feature-based ML & O(d) \text{ to } O(t) & Supervised training \\
SKADE & Ensemble model & $O(t)$ & Supervised training \\
SWI & Linear sequence index & $O(L)$ & None \\
Solupro & Feature-based ML & O(d) \text{ to } O(t) & Supervised training \\
EPSOL & Ensemble / ML & $O(t)$ & Supervised training \\
NetSolP & Deep neural network & $O(L \cdot k)$ & Supervised training \\
DeepSol-E & CNN-based model & $O(L \cdot k)$ & Supervised training \\
PLM\_Sol & Transformer (PLM-based) & $O(H \cdot L^2 \cdot h)$ & Pretrained + fine-tuned \\
\textbf{Composite-$\delta$} & Interpretable linear score & $\mathbf{O(L)}$ & None \\
\bottomrule
\end{tabular}
\end{adjustbox}
\label{tab:complexity_comparison}
\end{table}

For clarity, we define the notation used in the complexity analysis as follows: 
$L$ denotes the sequence length, 
$d$ the number of input features, 
$t$ the number of trees or ensemble components, 
$k$ the kernel or filter size in convolutional models, 
$H$ the number of transformer layers,
$h$ the number of attention heads per layer. 
All complexity estimates refer to inference-time computational cost under standard implementations.

The redundancy-aware composite-$\delta$ baseline achieved $\mathrm{AUC} = 0.6240$ and $\mathrm{MCC} = 0.1746$. Although this performance remains below that of high-capacity protein language model architectures such as PLM\_Sol \cite{zhang2024plmsol}, it is comparable to, and in several cases exceeds, traditional physicochemical feature-based predictors reported in the literature. Importantly, the composite-$\delta$ formulation involves no parameter fitting, no embedding extraction, and no hyperparameter optimization. The decision rule is fully determined by robust scaling \cite{Huber1981,Hampel1986} and statistically estimated effect sizes derived directly from empirical distributions. Consequently, the observed discrimination reflects separability of sequence-level biochemical features rather than model-driven representation learning.

From a computational standpoint, it is important to clarify that the computational complexity of the proposed composite-$\delta$ model depends on the level of representation considered. When applied directly to raw amino acid sequences, feature extraction requires a single pass over the sequence, resulting in a linear time complexity of $O(L)$, where $L$ is the sequence length. 

However, once sequence-derived features are computed, the composite-$\delta$ score itself is obtained through a simple linear combination, which operates in constant time $O(1)$. Therefore, the model is best interpreted as having $O(L)$ end-to-end complexity with an $O(1)$ scoring step.

This distinction is important in practice, as feature extraction is computationally inexpensive and scales linearly with sequence length, while the absence of training, parameter optimization, or iterative inference makes the overall framework substantially more efficient than high-capacity models such as protein language models, which typically scale at least quadratically with sequence length. In practical settings, where sequence-derived features are often precomputed or cached, the effective runtime of the scoring step becomes negligible. In contrast, classical machine-learning predictors based on handcrafted descriptors scale at least linearly with the number of features or ensemble components ($O(d)$–$O(t)$) \cite{bishop2006pattern}. Convolutional neural networks introduce sequence-length-dependent cost ($O(L \cdot k)$) \cite{lecun1998gradient}, while transformer-based protein language models incur quadratic complexity with respect to sequence length due to the self-attention mechanism ($O(H \cdot L^2 \cdot h)$) \cite{vaswani2017attention}. This progression corresponds to substantially higher computational cost.

The moderate performance gap between composite-$\delta$ and PLM-based architectures therefore reflects a clear trade-off between representational capacity and computational efficiency. While transformer models capture higher-order contextual interactions at substantial resource cost, the present results demonstrate that a low-dimensional, redundancy-controlled linear formulation retains a non-trivial portion of discriminative signal with negligible computational overhead.

From a mechanistic perspective, the achieved performance indicates that global physicochemical descriptors encode a measurable but limited solubility signal. The modest AUC and MCC values are consistent with the substantial distributional overlap observed in univariate analyses and reinforce the interpretation of solubility as a weak-signal, low-dimensional, and context-dependent phenotype \cite{chiti2006aggregation,chiti2017misfolding}. Rather than suggesting inadequacy of classical descriptors, these findings clarify their role: global sequence-derived features provide an interpretable empirical reference for the level of discrimination and reflect the interpretable physicochemical axes upon which more complex models may implicitly build. In this sense, the composite-$\delta$ baseline serves as both a transparent reference model and a mechanistic anchor for evaluating the added value of high-capacity predictive frameworks under explicit computational constraints. In addition to these mechanistic insights, it is important to consider the robustness of the proposed framework. To assess robustness, we note that the composite-$\delta$ formulation is derived directly from global distributional properties and does not involve optimization or parameter tuning beyond direct empirical estimation from the same dataset, the framework may be less sensitive to dataset partitioning, although this was not formally evaluated compared to trained models; formal stability assessment is left for future work. Future work may evaluate stability under resampling; however, given the large sample size ($N>78{,}000$), variance in estimated effect sizes is expected to be limited, although not formally evaluated.

\section*{Conclusion}

This study presents a large-scale, statistically controlled and interpretable analysis of sequence-derived biochemical determinants of protein solubility using a dataset of 78{,}031 labeled proteins. By integrating non-parametric testing, multiple-testing correction, effect size estimation, uncertainty quantification, ROC-based evaluation, and redundancy analysis, we explicitly distinguished statistical detectability from practical discriminative magnitude.

Our results show that soluble and insoluble proteins are consistently distinguishable at the distributional level across a wide range of descriptors. However, these differences are generally small in magnitude and exhibit substantial overlap between classes. This finding establishes that protein solubility at the sequence level operates in a weak-signal regime, in which no single descriptor provides strong standalone discrimination. Instead, separability arises from the coordinated contribution of multiple physicochemical features with modest individual effects.

Importantly, this analysis refines the interpretation of widely accepted solubility determinants. While factors such as sequence length, molecular weight, charge, and hydrophobicity have been extensively reported, our results demonstrate that their practical effect sizes are considerably smaller than often implied by statistical significance alone. By explicitly quantifying effect magnitude, this study distinguishes between features that are statistically detectable and those that contribute meaningfully to discrimination.

Redundancy analysis further reveals that much of the sequence-level signal is structured along a small number of latent physicochemical axes. Strong collinearity among size-related descriptors indicates that multiple high-ranking features reflect a shared structural burden dimension, whereas charge-related features represent a largely independent electrostatic axis. After redundancy filtering, a parsimonious two-dimensional composite based on sequence length and negative charge proportion retained measurable discriminative ability, supporting the interpretation that the analyzed feature space may be organized along a small number of dominant physicochemical axes.

These findings also inform the nature of weak-signal interactions. The observed patterns are consistent with predominantly additive contributions of global sequence-derived physicochemical features, although higher-order dependencies were not explicitly modeled here. While higher-order structural or contextual representations may further improve predictive performance, such approaches typically involve multi-stage inference from sequence to higher-level features, which can introduce cumulative error and substantially increase computational complexity. In contrast, the present framework isolates the intrinsic contribution of first-order sequence descriptors, providing a transparent baseline for understanding their standalone explanatory power.

An important limitation of the present framework is that it operates on global sequence-derived descriptors and therefore does not explicitly account for positional or contextual effects along the amino acid sequence. In biological systems, the functional role of a given residue is strongly dependent on its position and local sequence environment. Identical amino acids may contribute differently to folding stability, aggregation propensity, or solvent exposure depending on their structural context \cite{rives2021plm}. Such context-dependent effects are partially captured by high-capacity protein language models, which treat sequences as ordered and continuous objects and implicitly model higher-order dependencies. In contrast, the present study adopts a deliberately simplified representation in which sequences are treated as collections of compositional and global physicochemical features, thereby ignoring sequential ordering and positional information. This abstraction enables transparent interpretation and low computational complexity but inherently limits the ability to capture higher-order interactions. Consequently, the proposed framework should be understood as isolating the contribution of global, first-order physicochemical signals, providing a lower-bound characterization of sequence-based solubility information in the absence of contextual modeling. Future work may extend this framework by incorporating position-aware or context-sensitive representations while preserving statistical interpretability.

All dataset partitions were merged to maximize statistical power for descriptive distributional analysis. Accordingly, no claims regarding out-of-sample predictive performance are made. The composite-$\delta$ index should therefore be interpreted as a data-dependent statistical reference rather than as a trained predictive model.

Although the proposed composite-$\delta$ index achieves moderate descriptive discrimination compared to high-capacity models, it offers a favorable trade-off between interpretability, computational efficiency, and practical applicability. The model operates in linear time with respect to sequence length, with a constant-time scoring step, requires no training, and can be readily deployed in resource-constrained experimental environments where rapid pre-screening of protein solubility is needed. In such settings, a transparent and computationally inexpensive model can provide immediate value for preliminary screening, guiding experimental design before committing to more costly predictive pipelines.

Overall, this work establishes a statistically principled and interpretable reference framework for sequence-based solubility analysis. By quantifying effect size, redundancy, and low-dimensional structure in a large-scale dataset, this study provides a transparent baseline that clarifies the role of classical descriptors, refines their biological interpretation, and contextualizes the added value of more complex modeling approaches. The proposed composite-$\delta$ formulation should therefore be interpreted as an interpretable statistical reference that characterizes the magnitude, redundancy, and descriptive separability of global sequence-derived solubility signals. It is intended to provide a transparent point of comparison for evaluating the added value of more complex machine-learning and language-model-based approaches, rather than to serve as a competitive predictive model.

\section*{Supporting information}

% Include only the SI item label in the paragraph heading. Use the \nameref{label} command to cite SI items in the text.
Supplementary tables (Tables S1-S5) and figures (Figures S1-S4), containing detailed statistical results, model comparisons, and additional analyses, are provided as a separate Supporting Information file accompanying this manuscript.

\subsection*{Data and Code Availability}

The protein solubility dataset analyzed in this study was obtained from the benchmark dataset published by Zhang \textit{et al.}~\cite{zhang2024plmsol} and is publicly accessible via Zenodo at \url{https://zenodo.org/records/12881509}. 

All scripts used for sequence preprocessing, feature extraction, statistical testing, composite score construction, and figure generation were developed by the authors. The complete analysis pipeline, together with processed feature matrices, statistical summaries, composite score outputs, and manuscript figures, is publicly available at:

\texttt{https://github.com/huyhoang6723/protein-solubility-effectsize.git}

Within the repository, the file \texttt{data.zip} contains the dataset splits used in this study, while \texttt{results.zip} includes all derived features and analytical outputs reported in the manuscript. In addition, the full feature matrix used for all statistical analyses (including all 36 descriptors for each protein sequence) is provided in the repository (file: \texttt{features per protein.csv}), enabling complete reproduction of all reported results. Raw sequence datasets are referenced to their original public source in order to respect licensing constraints. This structure ensures full computational reproducibility of all reported results. A permanent archived release with DOI will be created upon acceptance.

\textbf{Repository Structure}

\dirtree{%
.1 protein-solubility-effectsize/.
.2 README.md.
.2 LICENSE.
.2 .gitignore.
.2 code.ipynb.
.2 data.zip.
.3 Train\_dataset.fasta.
.3 validation\_dataset.fasta.
.3 test\_dataset.fasta.
.2 results.zip.
.3 stats\_features.csv.
.3 summary\_global.csv.
.3 summary\_by\_class.csv.
.3 composite\_delta\_metrics.csv.
.3 composite\_delta\_predictions.csv.
.3 features\_per\_protein.csv.
.3 corr\_4features\_spearman.png.
.3 roc\_top4.png.
.3 distribution length.png.
.3 distribution negative.png.
}

\section*{Acknowledgments}

The authors thank the contributors of the publicly available protein solubility benchmark dataset used in this study and acknowledge the open source software community for providing essential computational tools that enabled large-scale statistical analysis. We are grateful to colleagues and reviewers for constructive discussions and critical feedback that improved the clarity and rigor of this work.

This research received no specific grant from any funding agency in the public, commercial, or not-for-profit sectors. The authors declare no competing interests.
\nolinenumbers

% Please compile your BiBTeX database using the "plos2025.bst" BibTeX style.
% This file is part of the current package.
% A sample BibTeX file is also included as "plos_bibtex_sample.bib".
%
% or
%
% Type in your references following Vancouver style and reference formatting instructions
% available at https://journals.plos.org/plosone/s/submission-guidelines#loc-references
% \begin{thebibliography}{}
% \bibitem{}
% Text
% \end{thebibliography}

\begin{thebibliography}{33}

\bibitem{baneyx1999expression}
Baneyx F. Recombinant protein expression in \textit{Escherichia coli}. Curr Opin Biotechnol. 1999 Oct;10(5):411--421.
doi: \texttt{\href{https://doi.org/10.1016/S0958-1669(99)00003-8}{10.1016/S0958-1669(99)00003-8}}.

\bibitem{chiti2017misfolding}
Chiti F, Dobson CM. Protein misfolding, amyloid formation, and human disease. Annu Rev Biochem. 2017;86:27--68.
doi: \texttt{\href{https://doi.org/10.1146/annurev-biochem-061516-045115}{10.1146/annurev-biochem-061516-045115}}.

\bibitem{rosano2014ecoli}
Rosano GL, Ceccarelli EA. Recombinant protein expression in \textit{Escherichia coli}: advances and challenges. Front Microbiol. 2014;5:172.
doi: \texttt{\href{https://doi.org/10.3389/fmicb.2014.00172}{10.3389/fmicb.2014.00172}}.

\bibitem{wilkinson1991solubility}
Wilkinson DL, Harrison RG. Predicting the solubility of recombinant proteins in \textit{Escherichia coli}. Biotechnology (N Y). 1991 May;9(5):443--448.
doi: \texttt{\href{https://doi.org/10.1038/nbt0591-443}{10.1038/nbt0591-443}}.

\bibitem{chiti2006aggregation}
Chiti F, Dobson CM. Protein aggregation in disease and biotechnology. Annu Rev Biochem. 2006;75:333--366.
doi: \texttt{\href{https://doi.org/10.1146/annurev.biochem.75.101304.123901}{10.1146/annurev.biochem.75.101304.123901}}.

\bibitem{idicula2005structure}
Idicula-Thomas S, Balaji PV. Understanding the relationship between the primary structure of proteins and its propensity to be soluble on overexpression in \textit{Escherichia coli}. Protein Sci. 2005 Feb;14(2):582--592.
doi: \texttt{\href{https://doi.org/10.1110/ps.041129005}{10.1110/ps.041129005}}.

\bibitem{ventura2005aggregation}
Ventura S. Sequence determinants of protein aggregation: Tools to increase protein solubility. Microb Cell Fact. 2005;4:11.
doi: \texttt{\href{https://doi.org/10.1186/1475-2859-4-11}{10.1186/1475-2859-4-11}}.

\bibitem{chou1974prediction}
Chou PY, Fasman GD. Prediction of protein conformation. Biochemistry. 1974 Jan;13(2):222--245.
doi: \texttt{\href{https://doi.org/10.1021/bi00699a002}{10.1021/bi00699a002}}.

\bibitem{kyte1982simple}
Kyte J, Doolittle RF. A simple method for displaying the hydropathic character of a protein. J Mol Biol. 1982 May;157(1):105--132.
doi: \texttt{\href{https://doi.org/10.1016/0022-2836(82)90515-0}{10.1016/0022-2836(82)90515-0}}.

\bibitem{wasserstein2019pvalue}
Wasserstein RL, Schirm AL, Lazar NA. Moving to a world beyond $p < 0.05$. Am Stat. 2019;73(Suppl 1):1--19.
doi: \texttt{\href{https://doi.org/10.1080/00031305.2019.1583913}{10.1080/00031305.2019.1583913}}.

\bibitem{khurana2018deepsol}
Khurana S, et al. DeepSol: A deep learning framework for sequence-based protein solubility prediction. Bioinformatics. 2018 Aug;34(15):2605--2613.
doi: \texttt{\href{https://doi.org/10.1093/bioinformatics/bty166}{10.1093/bioinformatics/bty166}}.

\bibitem{hon2021soluprot}
Hon J, et al. SoluProt: Prediction of soluble protein expression in \textit{Escherichia coli}. Bioinformatics. 2021 Jan;37(1):23--28.
doi: \texttt{\href{https://doi.org/10.1093/bioinformatics/btaa1102}{10.1093/bioinformatics/btaa1102}}.

\bibitem{thumuluri2022netsolp}
Thumuluri V, et al. NetSolP: Predicting protein solubility in \textit{Escherichia coli} using language models. Bioinformatics. 2022 Feb;38(4):941--946.
doi: \texttt{\href{https://doi.org/10.1093/bioinformatics/btab782}{10.1093/bioinformatics/btab782}}.

\bibitem{zhang2024plmsol}
Zhang X, et al. PLM\_Sol: Predicting protein solubility by benchmarking multiple protein language models with the updated \textit{Escherichia coli} protein solubility dataset. Brief Bioinform. 2024;25(5):bbae404.
doi: \texttt{\href{https://doi.org/10.1093/bib/bbae404}{10.1093/bib/bbae404}}.

\bibitem{dunker2001disorder}
Dunker AK, et al. Intrinsically disordered protein. J Mol Graph Model. 2001;19(1):26--59.
doi: \texttt{\href{https://doi.org/10.1016/S1093-3263(00)00138-8}{10.1016/S1093-3263(00)00138-8}}.

\bibitem{uversky2013intrinsic}
Uversky VN. Intrinsic disorder-based protein interactions and their modulators. Chem Rev. 2013 Sep;113(9):6844--6879.
doi: \texttt{\href{https://doi.org/10.1021/cr300363r}{10.1021/cr300363r}}.

\bibitem{mann1947test}
Mann HB, Whitney DR. On a test of whether one of two random variables is stochastically larger than the other. Ann Math Stat. 1947 Mar;18(1):50--60.
doi: \texttt{\href{https://doi.org/10.1214/aoms/1177730491}{10.1214/aoms/1177730491}}.

\bibitem{benjamini1995controlling}
Benjamini Y, Hochberg Y. Controlling the false discovery rate. J R Stat Soc B. 1995;57(1):289--300.
doi: \texttt{\href{https://doi.org/10.1111/j.2517-6161.1995.tb02031.x}{10.1111/j.2517-6161.1995.tb02031.x}}.

\bibitem{cliff1993dominance}
Cliff N. Dominance statistics. Psychol Bull. 1993 Nov;114(3):494--509.
doi: \texttt{\href{https://doi.org/10.1037/0033-2909.114.3.494}{10.1037/0033-2909.114.3.494}}.

\bibitem{hodges1963estimates}
Hodges JL, Lehmann EL. Estimates of location based on rank tests. Ann Math Stat. 1963 Jun;34(2):598--611.
doi: \texttt{\href{https://doi.org/10.1214/aoms/1177704172}{10.1214/aoms/1177704172}}.

\bibitem{efron1993bootstrap}
Efron B, Tibshirani RJ. \textit{An Introduction to the Bootstrap}. Chapman \& Hall; 1993.

\bibitem{hanley1982roc}
Hanley JA, McNeil BJ. ROC curve interpretation. Radiology. 1982 Apr;143(1):29--36.
doi: \texttt{\href{https://doi.org/10.1148/radiology.143.1.7063747}{10.1148/radiology.143.1.7063747}}.

\bibitem{youden1950index}
Youden WJ. Index for rating diagnostic tests. Cancer. 1950 Jan;3(1):32--35.
doi: \texttt{\href{https://doi.org/10.1002/1097-0142(1950)3:1<32::AID-CNCR2820030106>3.0.CO;2-3}{10.1002/1097-0142(1950)3:1<32::AID-CNCR2820030106>3.0.CO;2-3}}.

\bibitem{spearman1904}
Spearman C. Association measurement. Am J Psychol. 1904;15(1):72--101.
doi: \texttt{\href{https://doi.org/10.2307/1412159}{10.2307/1412159}}.

\bibitem{conover1999}
Conover WJ. \textit{Practical Nonparametric Statistics}. 3rd ed. Wiley; 1999.

\bibitem{Huber1981}
Huber PJ. \textit{Robust Statistics}. Wiley; 1981.

\bibitem{Hampel1986}
Hampel FR, et al. \textit{Robust Statistics}. Wiley; 1986.

\bibitem{kramer2012charge}
Kramer RM, et al. Increased negative surface charge correlates with solubility. Biophys J. 2012;102(8):1907--1915.
doi: \texttt{\href{https://doi.org/10.1016/j.bpj.2012.01.060}{10.1016/j.bpj.2012.01.060}}.

\bibitem{chan2013solubility}
Chan P, et al. Soluble expression correlates with lack of positive charge. J Mol Biol. 2013;425(8):1427--1435.
doi: \texttt{\href{https://doi.org/10.1016/j.jmb.2013.01.030}{10.1016/j.jmb.2013.01.030}}.

\bibitem{bishop2006pattern}
Bishop CM. \textit{Pattern Recognition and Machine Learning}. Springer; 2006.

\bibitem{lecun1998gradient}
LeCun Y, et al. Gradient-based learning applied to document recognition. Proc IEEE. 1998;86(11):2278--2324.
doi: \texttt{\href{https://doi.org/10.1109/5.726791}{10.1109/5.726791}}.

\bibitem{vaswani2017attention}
Vaswani A, et al. Attention Is All You Need. NeurIPS. 2017.
doi: \texttt{\href{https://doi.org/10.48550/arXiv.1706.03762}{10.48550/arXiv.1706.03762}}.

\bibitem{rives2021plm}
Rives A, et al. Biological structure from protein language models. PNAS. 2021;118(15):e2016239118.
doi: \texttt{\href{https://doi.org/10.1073/pnas.2016239118}{10.1073/pnas.2016239118}}.

\end{thebibliography}

\end{document}
