## Supplementary figures and images for "Large-Scale Statistical Dissection of Sequence-Derived Biochemical Features Distinguishing Soluble and Insoluble Proteins"

### Analytical workflow.png

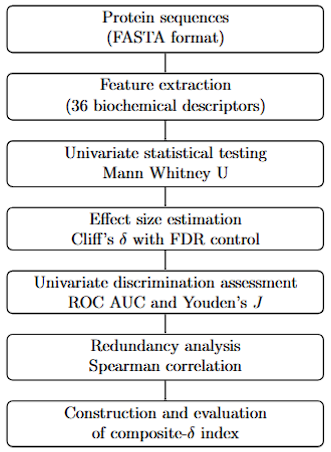

### corr_4features_spearman.png

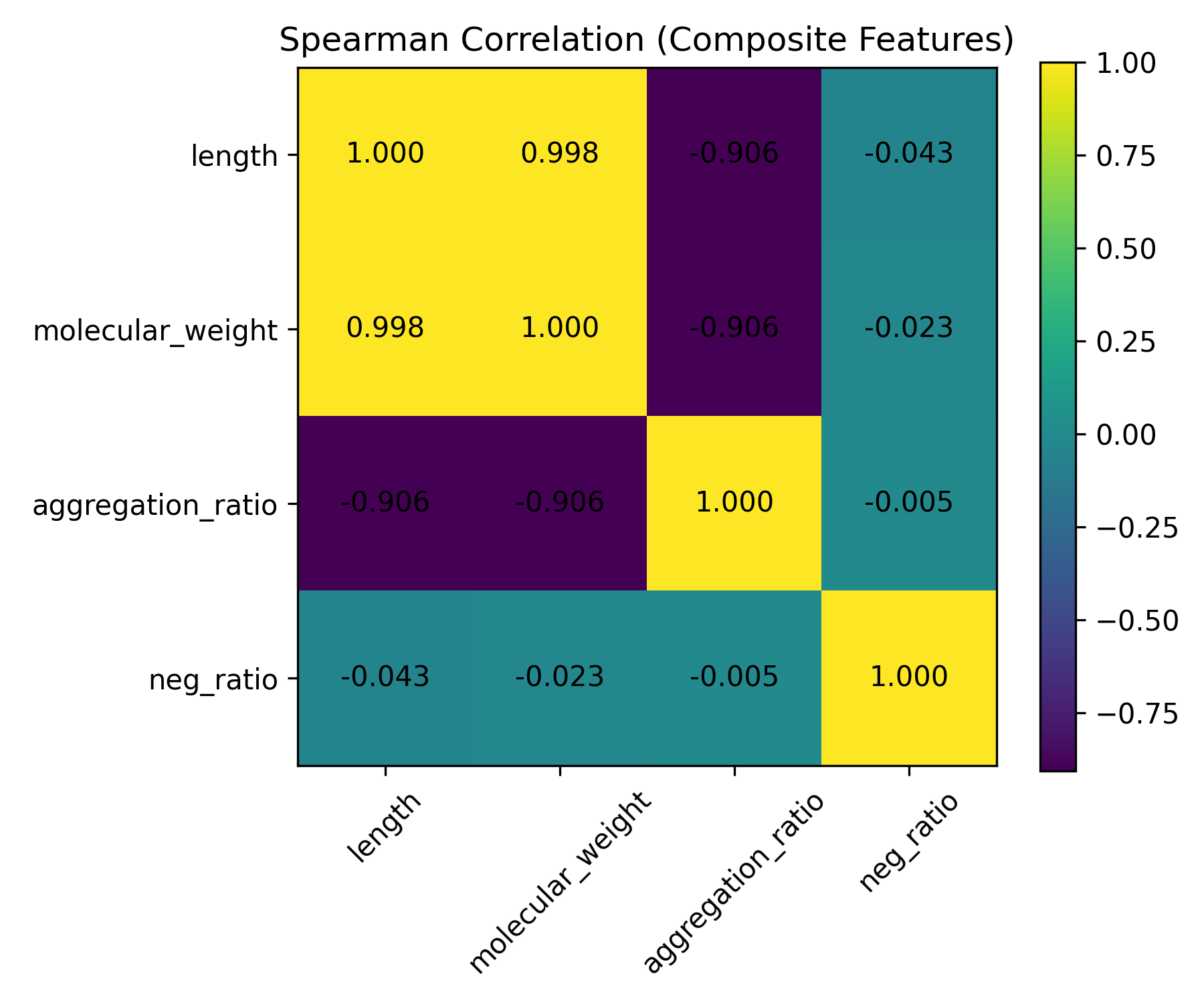

### distribution length.png

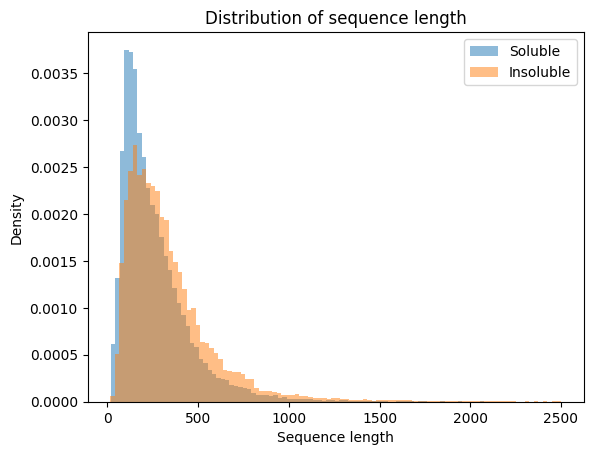

### distribution negative.png

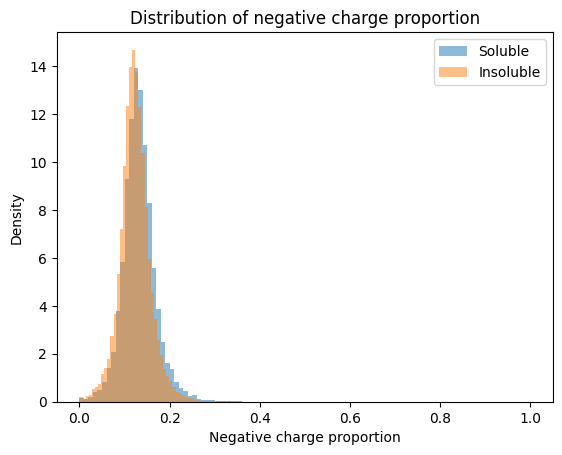

### roc_top4.png

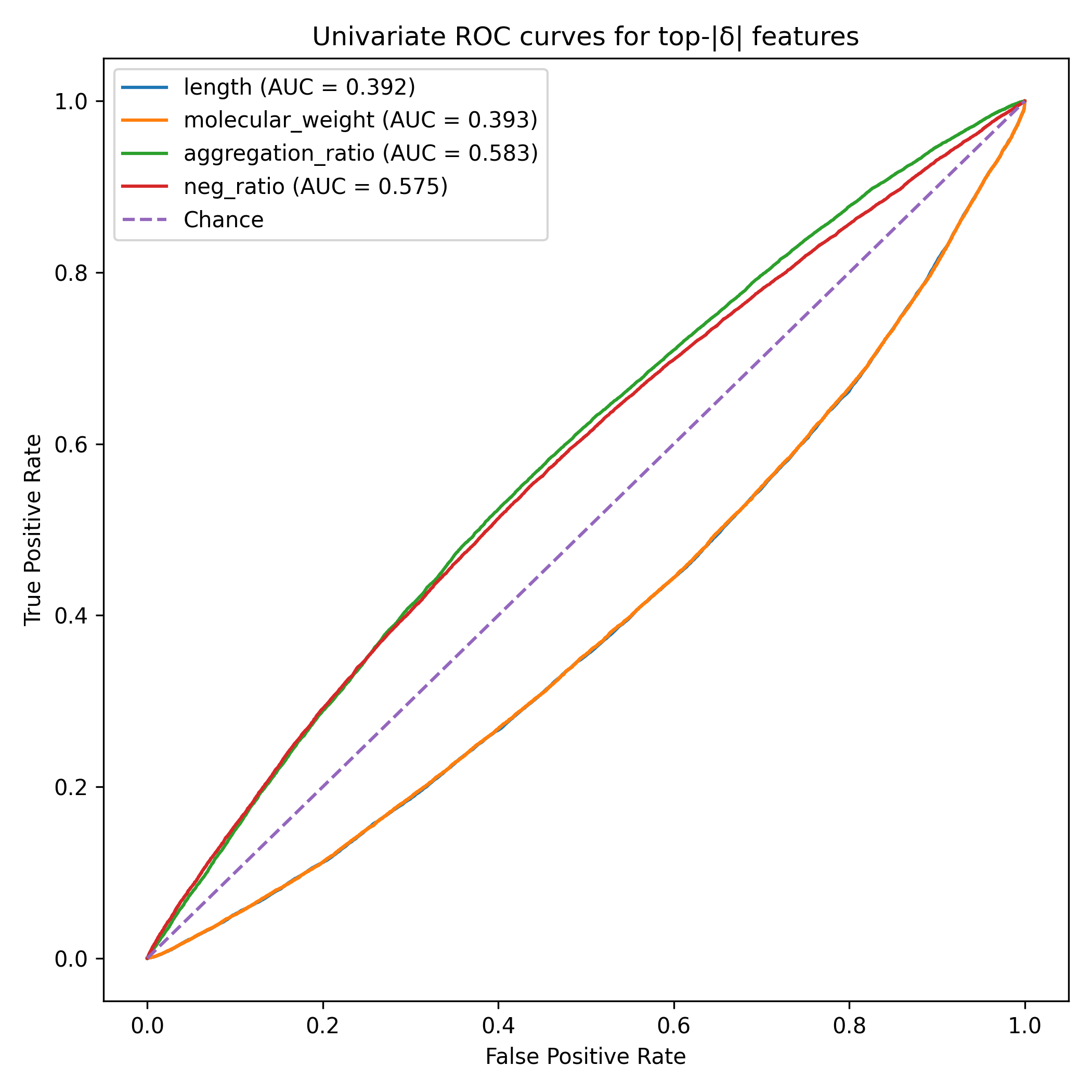
